# Resolving sulfation PTMs on a plant peptide hormone using nanopore sequencing

**DOI:** 10.1101/2024.05.08.593138

**Authors:** Xiuqi Chen, Jasper W. van de Sande, Justas Ritmejeris, Chenyu Wen, Henry Brinkerhoff, Andrew H. Laszlo, Bauke Albada, Cees Dekker

**Affiliations:** Department of Bionanoscience, Kavli Institute of Nanoscience Delft, Delft University of Technology, Delft, the Netherlands; Laboratory of Organic Chemistry, Wageningen University & Research, Wageningen, the Netherlands; Department of Physics, University of Washington, Seattle, WA, USA

## Abstract

Peptide phytohormones are decorated with post-translational modifications (PTMs) that are crucial for receptor recognition. Tyrosine sulfation on these hormones is essential for plant growth and development^1^. Measuring the occurrence and position of sulfotyrosine is, however, compromised by major technical challenges during isolation and detection^2^. We recently introduced a nanopore peptide sequencing method that sensitively detects PTMs at the single-molecule level^3^. By translocating PTM variants of the plant pentapeptide hormone phytosulfokine (PSK) through a nanopore, we here demonstrate accurate identification of sulfation and phosphorylation on the two tyrosine residues of PSK. Sulfation can be clearly detected and distinguished (>90%) from phosphorylation on the same residue. Moreover, the presence or absence of PTMs on the two close-by tyrosine residues can be accurately determined (>96% accuracy). Our findings demonstrate the extraordinary sensitivity of nanopore protein measurements, providing a new tool for identifying sulfation on peptide phytohormones and promising wider applications to identify protein PTMs.

---

Intercellular communication in plants is frequently regulated by secreted peptides, many of which contain at least one post-translational modification (PTM)^1,4^. Sulfation and phosphorylation on the tyrosine residue play critical roles in such signalling pathways and metabolism^5,6^. For peptide phytohormones such as phytosulfokine (PSK), plant peptide containing sulfated tyrosine (PSY), and root growth factor (RGF), the presence of a sulfate group on tyrosine is essential for their biological activity^7^. For example, the di-sulfated PSK is active at nanomolar concentrations only when both sulfate groups are present^8^. Recent bioinformatic estimates indicate that many more tyrosine residues can potentially be sulfated than currently known^9^. This suggests that sulfotyrosine residues tend to escape observation, likely due to the lability of the sulfoester bond during conventional mass spectrometry workflows, combined with biochemical purification protocols^2^ (Supp. Figure S1). Preserving and enriching sulfation PTM during sample preparation is far from optimized with known bias towards phosphotyrosine^2,10,11^, resulting in the underrepresentation of sulfotyrosine occurrence in the peptide phytohormone family. Importantly, the virtually identical masses of phosphotyrosine (79.966 Da modification) and sulfotyrosine (79.957 Da modification), differing by only 0.01 Da, make them difficult to distinguish by mass spectrometry. Furthermore, exact localization of the sulfation PTM is extremely challenging when multiple potential sites are encountered in a short fragment. This all calls for novel technologies that provide a high discriminatory power to differentiate between very similar PTMs and to locate the modification site.

Recent development in nanopore sequencing technology has provided new tools for identifying protein PTMs^12^. In this approach, a nanopore is inserted into the lipid bilayer and a target protein is linearized and slowly translocated through it. Amino acid residues within the pore constriction temporarily block the ionic current in slightly different ways, and variations in the signals correspond to the composition and sequence of the target protein. PTMs on proteins and peptides have recently been shown to produce distinct signals in nanopores^3,13,14^.

Here, we reveal how this novel nanopore sequencing technology can be used to distinguish between sulfotyrosine and the very similar phosphotyrosine within the PSK pentapeptide hormone consisting of the amino acid sequence YIYTQ. We find that single PTMs on PSK generate very distinct signals from the unmodified peptide and that sulfation can be accurately distinguished from phosphorylation. The exact location of the modified residue on the two potential PTM sites can also be clearly identified in all permutations at the single-molecule level. We thus show that nanopore sequencing offers a reliable, robust, and accessible method for determining PTMs on peptide phytohormones, with even single molecule sensitivity.

To enable the slow and step-wise translocation of the peptide through the MspA nanopore, one terminus of the peptide is covalently attached to a piece of single-stranded DNA (ssDNA) that is translocated through the pore by a Hel308 helicase (Figure 1A). On the other terminus, we added a negatively charged poly-aspartate (D_15_) tail to stretch the molecule under the applied voltage and to improve the efficiency of inserting the peptide into the pore. We synthesized the nine possible sulfation/phosphorylation PSK variants, *i*.*e*., no PTM, single PTM on either tyrosine (sulfation or phosphorylation), and PTMs on both residues. These were synthesized on a fifteen-aspartate chain by a solid-phase peptide synthesis protocol described previously^15^. The resulting peptides carry an azide group at the N terminus to enable strain-promoted alkyne-azide click attachment to a BCN-functionalized ssDNA (see Methods and Extended Figure 1-1 for the molecular structure). By applying a constant voltage bias across the lipid bilayer, the ionic current through the MspA nanopore was recorded and the translocation of the conjugate molecule was identified by the current blockade. The step-wise translocation induced by the Hel308 motor protein produced signal steps in the ionic current, as described previously^16,17^ (Figure 1C). With established signal prediction for DNA sequencing^18^, the onset of the peptide signals could be identified as following the aligned DNA signal steps (Figure 1D).

**Figure 1.**
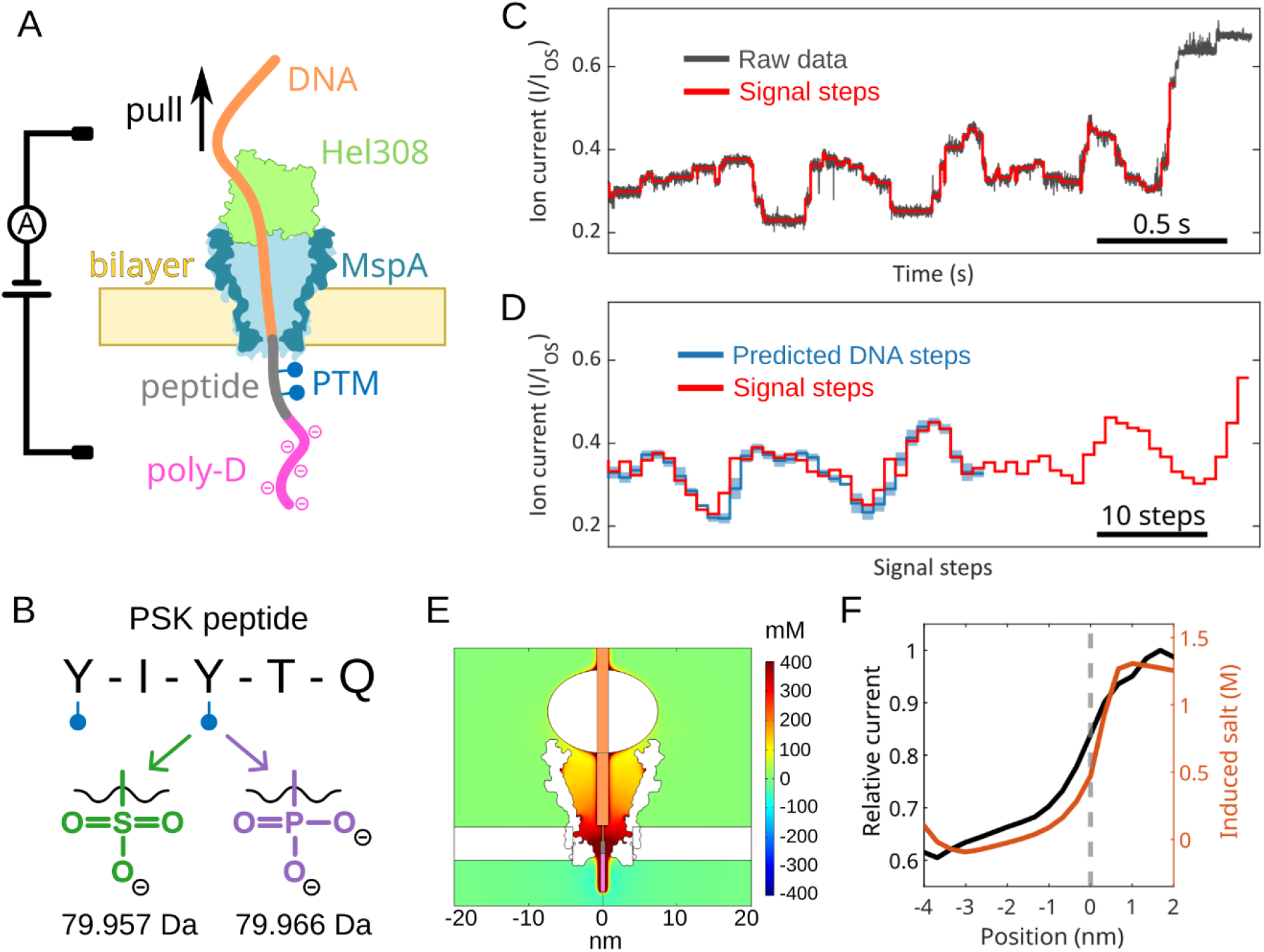
Detecting post-translational modifications on a PSK peptide with nanopore sequencing. (**A**) A DNA-peptide conjugate molecule translocates a MspA nanopore under a voltage bias, until the Hel308 that is bound to the DNA is stuck at the top of the pore. The Hel308 motor protein slowly pulls the DNA upward, generating a step-wise ionic current as the DNA-peptide conjugate passes the narrow pore constriction. (**B**) Two tyrosine (Y) residues on the pentapeptide PSK can be modified by either sulfation or phosphorylation, carrying one or two negative charges, respectively. Masses are calculated in their protonated forms. (**C)** Example ionic current trace from the double sulfation sample. The open-state current (I_OS_) of the nanopore is used to normalize the current blockades across different translocation events. Step-like signals are identified and used to characterize the analyte. (**D**) Given the known DNA sequence, the predicted DNA signals are used to align and segment the full-length signals from the DNA-peptide conjugate molecule. The enzyme occasionally skips and back-steps, creating alignment shifts. (**E**) COMSOL simulation demonstrates the elevation of local salt concentrations near the pore mouth, induced by the densely charged poly-D tail (the central rod models, from top to bottom, the DNA, linker, peptide, and poly-D tail). Colour bar on the right denotes the scale of additional salt compared to the normal salt concentration of 400 mM. (**F**) Correlation of induced additional salt concentration at the nanopore constriction and ionic current during translocation. Dashed line marks the first residue from poly-D tail at the pore constriction.

For all peptide variants in this study, we observed a significant ionic current increase near the end of the translocation event, which served as a consistent reference for thresholding the end of the peptide signals (Extended Figure 1-2). Finite-element analysis with COMSOL Multiphysics (see Figure 1E and Methods) showed that this signal ramp can be attributed to the ion enrichment in the nanopore due to the dense negative charges on the poly-D tail. Known as ionic concentration polarization^19^, the charges on the peptide raise the local ion concentration and hence increase the ionic current when this part of the peptide is located near the nanopore constriction (Figure 1F). The onset of this current increase occurs slightly before the tail starts translocating the nanopore (Figure 1F, dashed line). A recent molecular dynamics (MD) study discussed a similar effect from charged polymers during translocation^20^. A less densely-charged poly-T ssDNA tail resulted in only a minor ionic current elevation (Supp. Figure S2).

We find that we can robustly distinguish signal traces from different PTM variants of the PSK hormone. Upon collecting many single-molecule events (∼100 traces for each sample), a consensus signal was constructed for each PTM variant through an improved Hidden Markov Model solver based on previous works^3,16^. The new implementation here incorporated contamination tolerance of bad signal levels, which happen stochastically during measurement (see Supplementary Methods). The different PTM states of the YIYTQ pentapeptide yielded significantly different consensus traces (Figure 2A). Phosphorylation on the 2^nd^ tyrosine induced a pronounced signal peak in the middle of the consensus, similar to previous observations in phosphorylated immunopeptides^3^. Sulfation on the same tyrosine residue similarly produced a signal peak but with a lower amplitude and an earlier onset when compared to phosphorylation (Figure 2A). This result is consistent with local salt modulation and charge-induced stretching of the polymer^3,20^. The di-anionic phosphotyrosine versus mono-anionic sulfotyrosine induces a higher local salt concentration and generates stronger stretching, and accordingly a delayed and higher signal peak. These distinct patterns from the sulfation and phosphorylation consensus resulted in highly accurate (>90%) variant identification of individual reads (Figure 2B, see Methods). Only a handful of the phosphorylated molecules were mistaken for sulfated molecules, mostly due to finite measurement noise. In contrast to the effects of the PTM on the 2^nd^ tyrosine, peptide variants with only the modified 1^st^ tyrosine showed less pronounced differences (Supp. Figure S3), which nevertheless could be mutually well-distinguished (91%) in variant calling.

**Figure 2.**
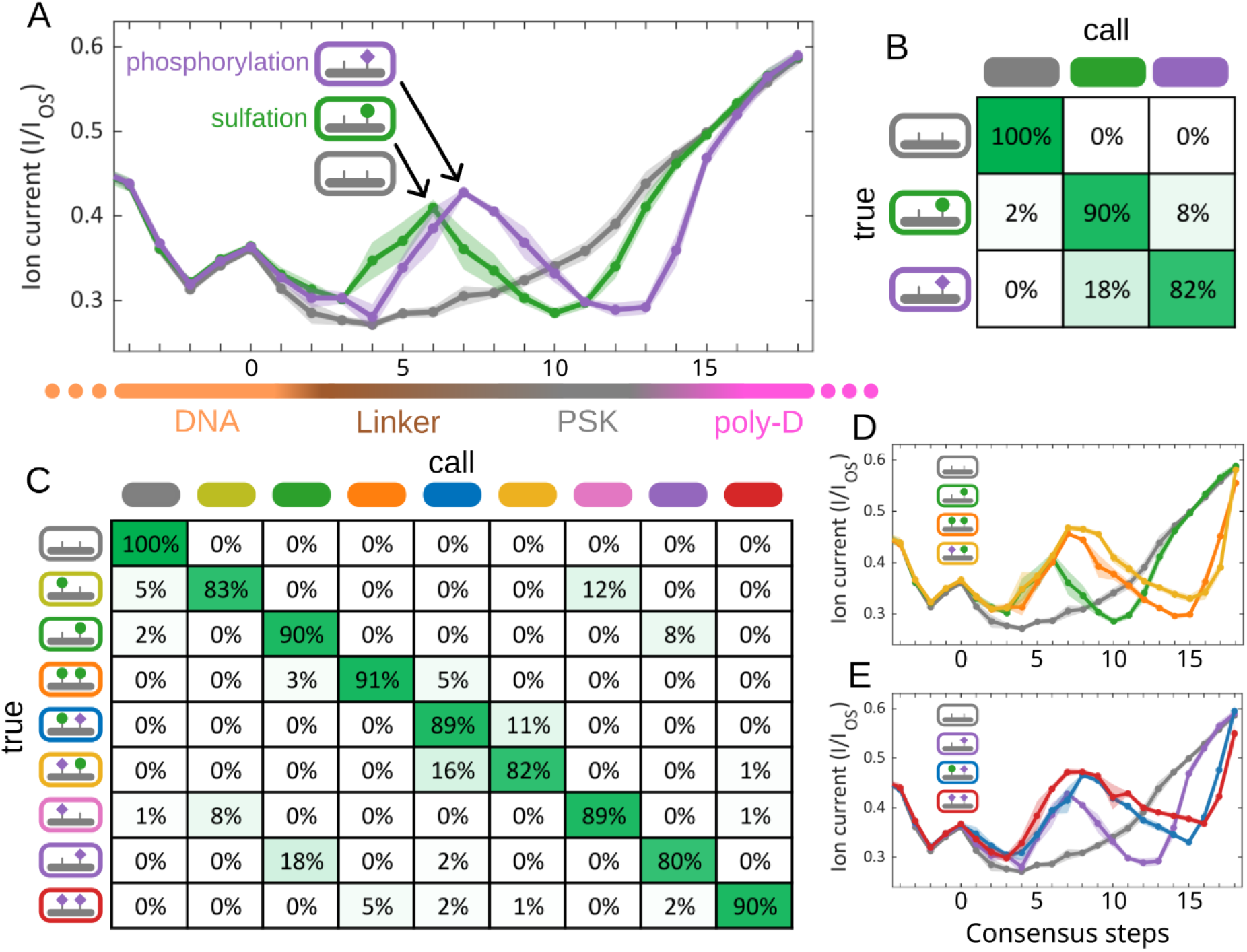
Sulfation and phosphorylation PTMs on the PSK hormone can be accurately identified. (**A**) Signal consensus traces from the unmodified PSK pentapeptide (grey), single sulfation (green), and single phosphorylation (purple). The starting DNA signals (orange region at the left) are identical across variants. Signal variation starts when the linker approaches the constriction site. The peptide signal is heavily influenced by the negatively charged poly-D tail (pink) that displays the signature ionic current ramp at the end of the traces. Means and standard deviations of the consensus steps are plotted as dots and shaded area. (**B**) Confusion matrix for three variants of panel A. Rows describe the true labels and columns describe the call results. (**C**) Same as B but for all variants. From top to bottom, n = 96, 130, 97, 115, 89, 134, 150, 132, 175. (**D** and **E**) Cumulative effects of additional PTMs on both tyrosine residues result in higher ionic current, either when the 2^nd^ tyrosine is sulfated (D) or phosphorylated (E).

Expanding variant identification across all possible PTM permutations highlighted the robustness of our nanopore detection method. Samples from the double, single, or unmodified variants could be well distinguished with a consistent high accuracy varying between 82% and 100% (Figure 2C). As the negative charge on the peptide increases, either by adding another PTM or by switching from sulfation to phosphorylation, a higher ionic current was obtained (Figure 2 D and E, Extended Figure 2-1). In particular, when both tyrosine residues were modified, the four permutations of phosphorylation and sulfation revealed a narrow region in the consensus (Grey shade, Extended Figure 2-2) where the rising negative charges caused increasing signals. Because ionic current through MspA is most sensitive to charges within the constriction, we infer that locations with the largest current difference in the consensus correspond to steps where modified tyrosine residues are within the pore constriction. Control experiments with a longer linker length between the DNA and peptide created, as expected, a shifted tail pattern that identified the same region for peptide translocation (Cyan shade, Extended Figure 2-2).

Furthermore, we found that it is possible to accurately identify the location of the PTM within the peptide, *i*.*e*., whether the same PTM occurred on the first or on the second tyrosine residue on the PSK. As Figure 3A shows, the four permutations of the pentapeptide carrying sulfation at two tyrosine residues generated distinct consensus patterns, yielding excellent identification accuracies of 95% to 100% (Figure 3B). Two seemingly independent observations can be made from the respective tyrosine measurements. For the modified 1^st^ tyrosine, the traces did not show an early signal peak, but instead presented a delayed and steeper signal ramp at the end. For the modified 2^nd^ tyrosine, an early signal peak appeared, but the tail ramp occurred later than the unmodified pentapeptide, gradually converging with the unmodified peptide. Double modification on the peptide combined these effects of the two sites, with a slightly higher peak amplitude from the additional negative charge. The phosphorylation variants of the peptide consistently followed the same rules, with stronger amplitudes (Figure 3C-D), consistent with the higher charges from the phosphorylation PTM. Because of the distinct effects from the two tyrosine PTMs, the modified variants carrying two negative charges, either from one phosphate or two sulfate groups, can also be accurately identified (Extended Figure 3-1).

**Figure 3.**
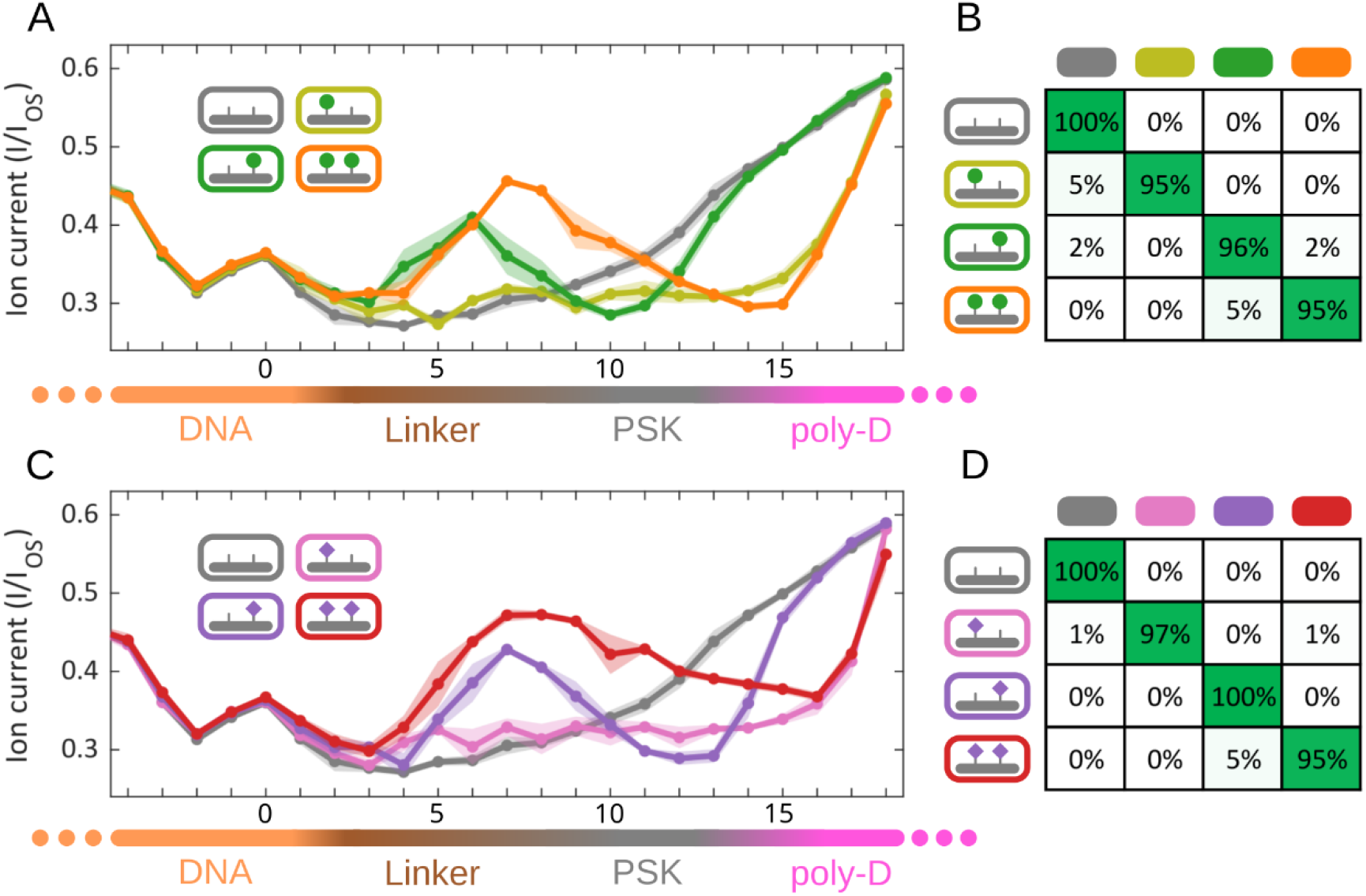
PTMs on two close-by tyrosine residues are well distinguished. (**A**) Consensus traces for the sulfation variants. The early signal peaks are related to the 2^nd^ tyrosine PTM while the tail pattern is related to the 1^st^ tyrosine PTM. (**B**) Confusion matrix of the sulfation variants. (**C**) Consensus traces for the phosphorylation variants. (**D**) Confusion matrix of the phosphorylation variants.

While we thus establish excellent discrimination between sulfation and phosphorylation PTMs on PSK, the molecular determinants that underly the peptide translocations are not *a priori* evident. For example, why does the signal peak from the modified 2^nd^ tyrosine appear so early in the linker region? Upon inserting a piece of PEG8 between the linker and PSK pentapeptide, the permutations of sulfation in the PEG8 variants conformed to the same patterns as observed before, *i*.*e*., early peak from the modified 2^nd^ tyrosine, and delayed tail from the modified 1^st^ tyrosine (Figure 3E). MD simulation indicated that the modified 2^nd^ tyrosine has a higher propensity to initiate a transient charge interaction with the positively charged arginine residues at the bottom of the MspA (Supp. Figure S4). This suggests that the early signal peak is not related to the linker, but more likely due to the closer proximity of the 2^nd^ tyrosine to the negatively charged poly-D tail. Recent nanopore studies also described ionic current alterations from charge interactions near nanopore constriction site^21,22^. By contrast, the delayed tail signals may be explained by hydrophobic interactions between the unmodified 1^st^ tyrosine with the pore inner surface (similar to observations described in a previous study^16^) that are disrupted by charged PTMs on this tyrosine. Charged PTMs on the peptide thus impact two distinct interactions at separate translocations steps (Extended Figure 3-3).

In this study, we applied single-molecule nanopore sequencing to detect and distinguish post-translational modifications with an isobaric mass on the plant peptide hormone phytosulfokine (PSK). While functional plant peptide hormones often carry sulfated tyrosine residues and act at extremely low concentrations in plants^7^, instability of tyrosine sulfation in typical mass spectrometry workflows has hindered investigation on this important family of peptides, especially when it comes to determining the site of modification^2^. We demonstrated that single-molecule nanopore measurements can be done with mild sample preparation conditions, and enable very accurate determination of sulfated and phosphorylated sites on peptides. Single sulfation or phosphorylation PTM on the peptide generated clearly distinguishable signal patterns. Permutations of PTMs on the two tyrosine residues revealed a surprising pattern where modifications on the 2^nd^ tyrosine gave rise to more pronounced signal changes. The two distinct effects from the PTMs on the respective tyrosine residues allowed for very accurate variant identification of these proximal modifications. The single-read accuracies range from 80 to 100%, which can be further improved to basically 100% by re-reading the same individual peptide multiple times by adjusting the experimental conditions^3^.

More generally, the current findings show the strength of nanopore sequencing technologies for PTM detection, especially for peptide phytohormones. Variant detection at the highest distinguishing power is readily attainable, as evidenced in our work where we demonstrated the extraordinary distinguishing power between even two closely positioned and very similar PTMs as sulfation and phosphorylation. With a generic peptide conjugation strategy for DNA attachment available^23^, our nanopore methodology can be widely applied to native plant peptide samples. Challenging post-translational modifications can be sensitively detected once measurement references are established, which aids in elucidating the complex peptide-based communication networks in ecosystems.

## Methods

### Solid-phase peptide synthesis of PSK-like peptides

Amino acids and peptide synthesis reagents were purchased from Novabiochem. Modified amino acids Fmoc-Tyr(SO_2_ONp)-OH and Fmoc-Tyr(PO(OBzl)OH)-OH were purchased from Merck Life Science. Azidoacetic acid was purchased from TCI Europe. Azido-PEG_8_-NHS ester was purchased from Broadpharm. Peptides were synthesized following standard Fmoc/*t*Bu solid-phase peptide synthesis (SPPS) strategy in a split method^15^ (see Supplementary Information). Peptides were cleaved from the resin with different trifluoroacetic acid (TFA) cocktails depending on the N-terminal modification. Treatment of peptide with 2M NH_4_OAc at 45 °C for 40 h resulted in neopentyl removal from the sulfated tyrosine residues. Obtained deprotected peptides were purified by preparative HPLC and analysed with UHPLC-MS for quality control (Supp. Figures S6-S18).

### SPAAC peptide-oligonucleotide conjugation

BCN-modified DNA was custom synthesized and purchased from Thermo Fisher Life Sciences (USA) with UHPLC-MS quality control performed in-house (Supp. Figure S5). 1.3 mM of desired peptide (20 nmol) and 0.3 mM of BCN-DNA (5 nmol) were added to a 0.5 mL microcentrifuge tube and reacted in Mili-Q water overnight at room temperature. Samples were purified with Amicon® ultra-spin filtration units with 10 kDa molecular weight cut-off (MWCO) using phosphate buffer (50 mM sodium phosphate, pH 6.0). Obtained samples were analysed with RP-UHPLC-MS (Supp. Figures S6-S18). Concentrations were measured with a Nanodrop spectrophotometer and corrected for the extinction coefficient of the template DNA as determined by the supplier. Samples were aliquoted, snap-frozen, and subsequently lyophilized.

### Nanopore measurements

Nanopore experiments on DNA-peptide conjugate molecules were performed as described in previous studies^3,16^. DPhPC lipids purchased from Avanti Polar Lipids (SKU: 850356C). M2-MspA mutant was purified by Genscript. Hel308 used in this study is from *Thermococcus gammatolerans* (accession number WP_015858487.1) and was purified in-house. Teflon aperture on custom U-tube devices were painted with DPhPC lipids to form bilayers submerged in buffer H (400 mM KCl, 10 mM HEPES, pH 8.00). Cross-membrane voltage was set to 180 mV under the control of a National Instruments X series DAQ and operated with custom LabVIEW software. Around 0.5 μL (1 μg/mL) of MspA was added to the *cis* well until a signature 140-150 pA ionic current increase was observed, indicating single nanopore insertion. The *cis* well is then perfused with buffer H supplemented with 1 mM ATP and 10 mM MgCl_2_. Hel308 and conjugate molecules were added to the *cis* well to a final concentration around 100 nM and 10 nM, respectively. At 37 °C, ionic current data were acquired at 50 kHz sampling frequency using an Axopatch 200B patch clamp amplifier and filtered with 10 kHz 4-pole Bessel filter.

### Data processing of single-molecule events

Ionic current recordings were first down-sampled to 5 kHz, and translocation events were identified by thresholding the current with the open pore current. The open-state current (I_OS_) of the nanopore is used to normalize the current blockades across different translocation events. The events of conjugate molecules, containing both DNA and peptide signals, were selected by eye. The peptide signal section from each event was marked consistently at a defined location by referencing the predicted DNA signal pattern, derived from an 6-mer map^18^. Signal steps generated by the helicase movement were extracted by a change point algorithm, described previously^18^. These ionic current steps were filtered by excluding any levels outside the bounds of expected current values (I/I_OS_ < 0.15 or I/I_OS_ > 0.7) before being linearly calibrated by aligning to a fixed DNA prediction reference (similar to Figure 1D). The calibrated signal steps from the peptide were then trimmed at the end by thresholding the tail ramp at 0.6 relative current.

### Consensus generation and variant calling

The events from each peptide variant were randomly reorganized into two equal groups, one for consensus generation and one for variant calling. Signal steps in the peptide region were manually picked in a selection of typical events (n = 4–6) to approximate the step-wise current levels generated by the helicase without back-stepping or skipping. This process produced some good input guesses for the Hidden Markov model solver for the peptide consensus. Along with the peptide events in the consensus generation group, the input guess was used in a custom Baum–Welch script to solve the hidden Markov model using a maximum-a posteriori likelihood (MAP) algorithm (See Supplementary Information). This assigns likelihood values that each of the signal steps in the event were produced by a particular true template position within the constriction (helicase step number)^16^. The resulting model is the consensus for the peptide variant, with mean and standard deviation values for each helicase step. For the peptide events in the variant calling group, each event was aligned to all the consensus from peptide variants for an alignment likelihood. The consensus alignment with the highest likelihood was designated as the call label. The accuracy of variant call was the percentage of call labels matching the true labels of the peptide events.

### COMSOL simulation

Numerical simulations of MspA-helicase-peptide system was implemented in COMSOL Multiphysics 5.4 with a two-dimensional axial symmetrical domain. The simulation included the fluid domain, the lipid membrane, MspA nanopore, helicase, and DNA-linker-peptide strand. With the contour of M2-MspA nanopore (PDB: 1UUN), the charged residues in the inner wall of the nanopore protein were marked at positions of 63 (−), 118 (+), and 134 (+). The DNA-linker-peptide strand was approximated with cylindrical columns, with corresponding thickness. The single-strand DNA (ssDNA) carried −1e/nucleotide, the linker was neutral in charge, and the D_15_ tail carried −1e/amino acid. The charge of tyrosine residues was set at 0, −1e, and −2e, for unmodified, sulfated, and phosphorylated states. To calculate the ionic current, ion flux was integrated on the cross-sectional area located at narrowest restriction of the nanopore. The relative current is based on the highest current during translocation. See Supplementary Information for detailed descriptions.

## Acknowledgments

The authors acknowledge E. van der Sluis and A. van den Berg for Hel308 purification, and A. Barth and J. van der Torre for helpful discussions. This work was supported by funding from the Dutch Research Council (NWO) project NWO-I680 (SMPS) (C.D.); European Research Council Advanced Grant 883684 (C.D.); and NIH NHGRI project HG012544-01 (C.D.) and from Wageningen University & Research (B.A. and J.W.v.d.S.).

## Data and code availability

The raw data and the code for analysis will be deposited in Zenodo.

## Author contributions

C. D. and B. A. conceived the study. X. C. performed the nanopore data analysis and drafted the manuscript. J. W. v.d. S. synthesized and analysed all DNA-PSK constructs. J. R. performed the nanopore measurements. C. W. performed COMSOL simulation. H. B. and A. L. developed the new alignment and consensus generation script. B. A. performed the MD simulations. All authors participated in manuscript revision.

## Extended figures

**Extended Figure 1-1.**
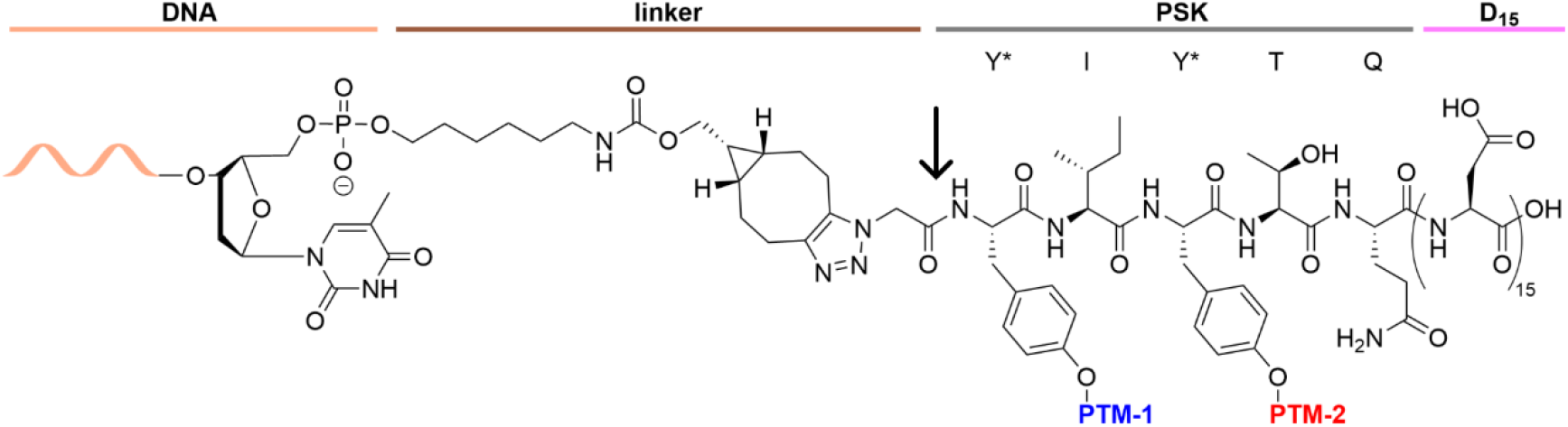
The chemical structure of the linker connecting the DNA and peptide. The 5’ end of the DNA is modified and conjugated with the N terminus of the PSK through click chemistry. The C terminal of the peptide is composed of fifteen aspartate (D_15_). The arrow indicates the PEG8 insertion site for some samples.

**Extended Figure 1_2.**
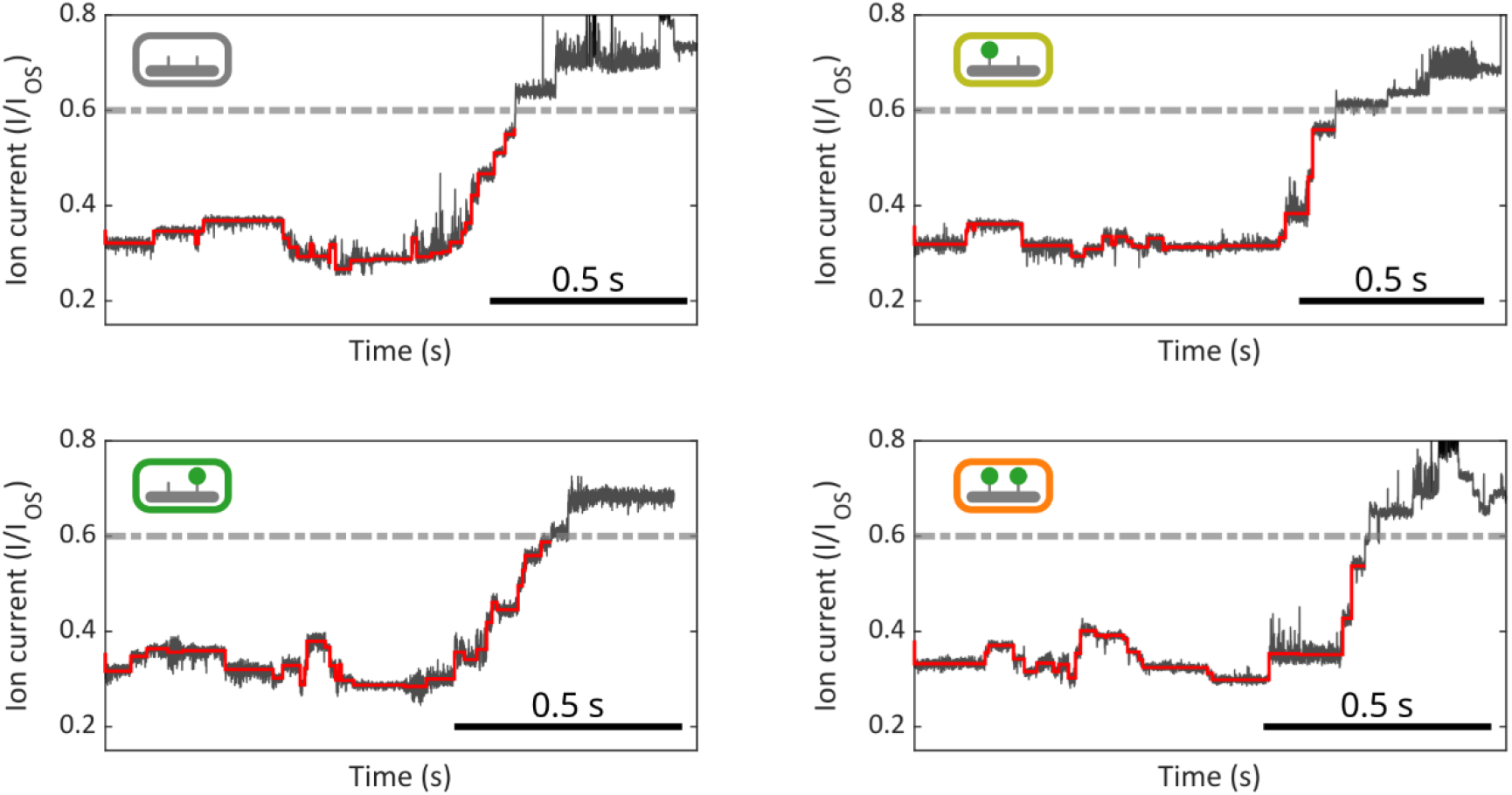
Example traces showing raw data that are thresholded at the poly-D tail. Above a certain limit (relative current at 0.6 in the dashed line), the nanopore measurement of peptide variants become more noisy and less consistent, likely due to fluctuations when the peptide is about to exit the nanopore. Consequently, all single-molecule events in this study are thresholded at I/I_OS_ = 0.6 at the ramp for reliable analysis.

**Extended Figure 2_1.**
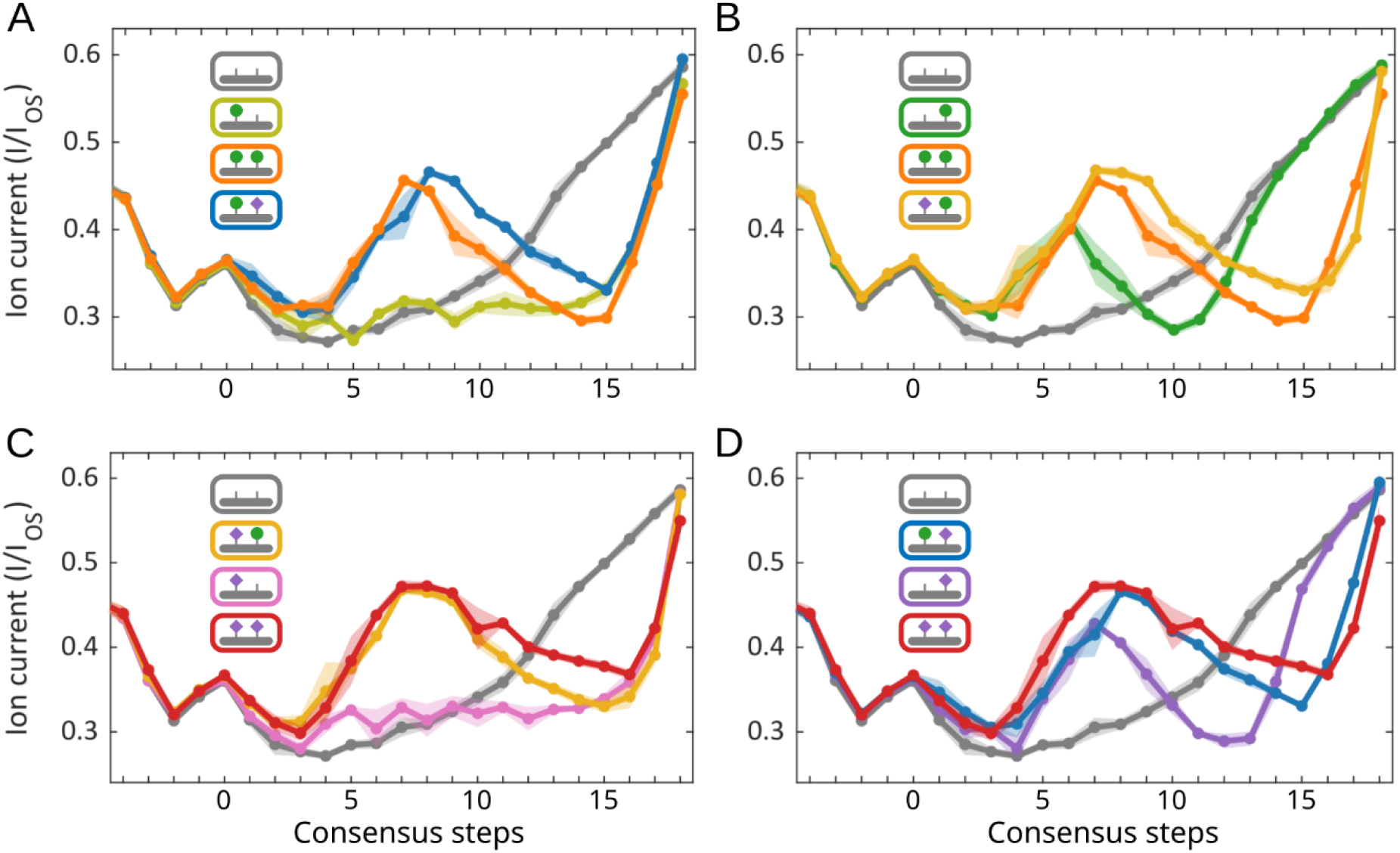
Site-specific comparisons of PTM variants. Signal consensus of variants carrying sulfation on 1^st^ tyrosine (**A**), sulfation on 2^nd^ tyrosine (**B**), phosphorylation on 1^st^ tyrosine (**C**), phosphorylation on 2^nd^ tyrosine (**D**). The unmodified peptide (grey) is used as a reference trace in all plots. Means and standard deviations of the consensus steps are plotted as dots and shaded area, respectively.

**Extended Figure 2_2.**
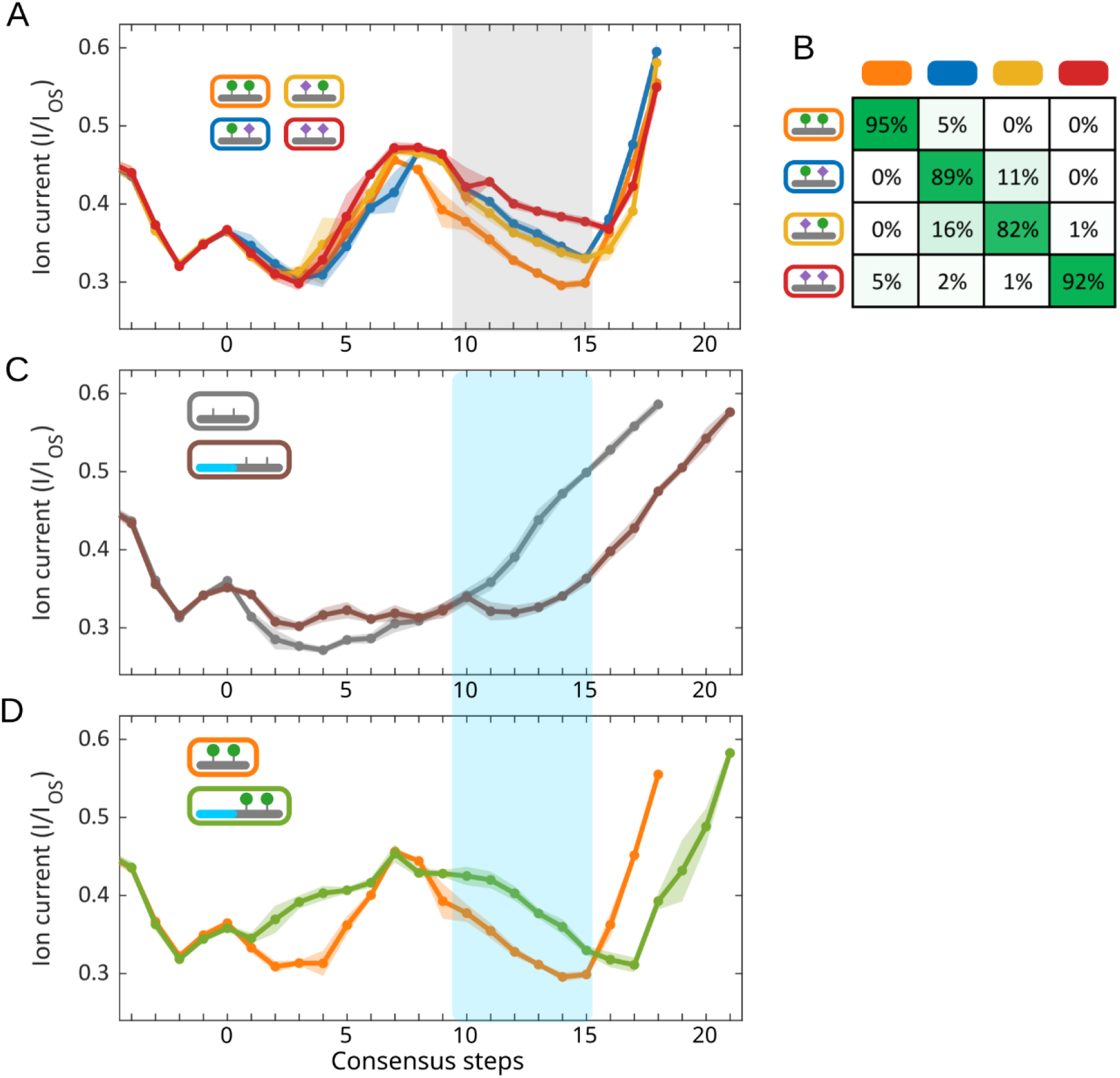
Double-modification variants and PEG linker variants locate PSK translocation steps. (**A**) Double modification variants highlighted a region with the most signal differences near the modified PSK. (**B**) Confusion matrix of the double modification variants. (**C** and **D**) The PEG8 linker insertion clearly shifted the tail signal pattern in the non-modified and double modified variants, likely reflecting the PEG8 insertion site. The blue shaded region corresponds well to the grey shaded region, positioning the translocation of the PSK peptide.

**Extended Figure 3-1.**
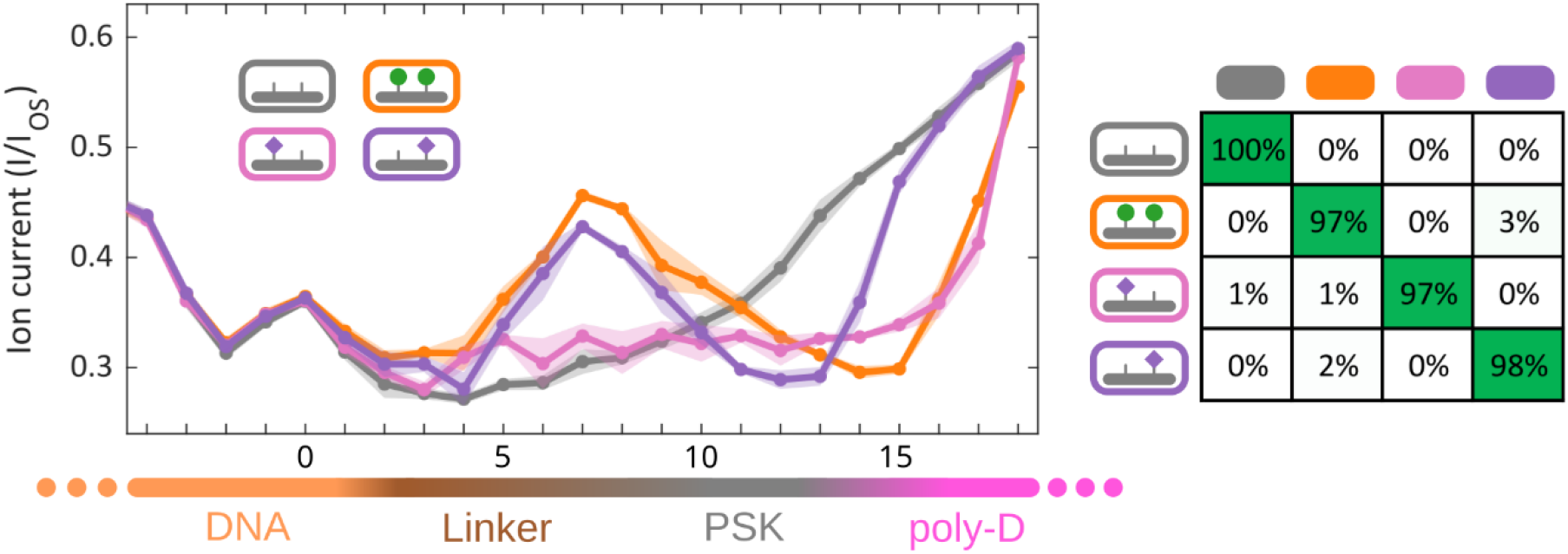
Consensus comparison and variant calling of peptides carrying two negative charges. The peptide variant either carries one phosphate group (pink or purple) or two sulfate group (orange).

**Extended Figure 3-2.**
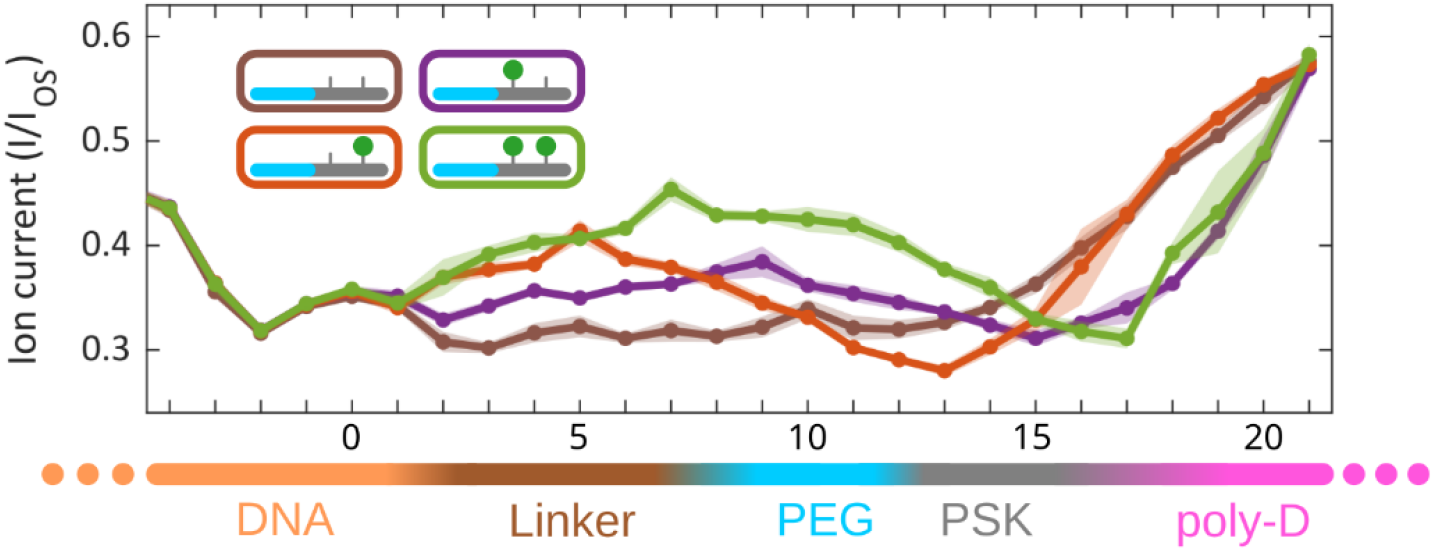
Consensus traces for the sulfation variants with an inserted PEG8 linker. The PEG8 insertion shifted the tail ramps by a few steps while tail patterns were still governed by the 1^st^ tyrosine PTMs. The early signal peaks were also maintained by 2^nd^ tyrosine PTMs.

**Extended Figure 3-3.**
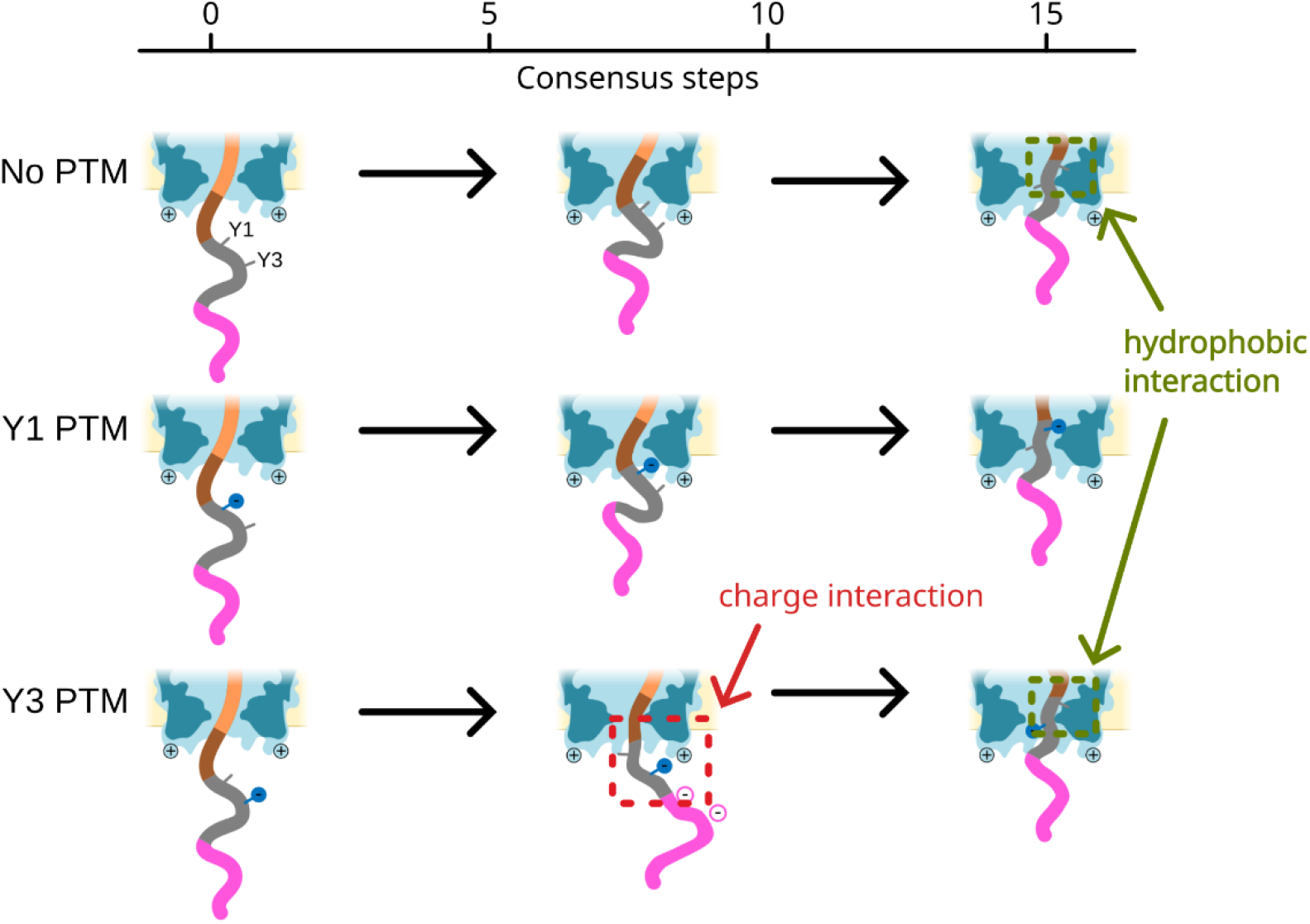
Possible molecular conformations during translocation. Early during translocation, charged modifications on the 2^nd^ tyrosine (Y3) interact with the positively charged residues at the bottom of MspA, producing a signal peak. This interaction is unique to Y3 likely due to the proximity to the densely charged poly-D tail. After the PSK enters the nanopore constriction, the unmodified 1^st^ tyrosine (Y1) interacts with the pore inner surface and increases the ionic current. Charged modifications on Y1 disrupts this interaction and cause the tail pattern to shift.

