## SupplementaryInformation for "Resolving sulfation PTMs on a plant peptide hormone using nanopore sequencing"

SI Methods

- Ultra-high performance liquid chromatography coupled to MS (RP-UHPLC-MS)
- General procedure for Solid-Phase Peptide Synthesis of PSK-like peptides
- COMSOL simulation on peptide translocation
- Brief overview of the HMM solver with contamination tolerance
- Molecular dynamics simulation

SI Tables

- Table 1. Sequences of DNA and DNA-peptide conjugates in this study

SI Figures

- S1. CID results on sulfation under mass spectrometry
- S2. COMSOL simulation on charge-induced salt concentration and ionic current elevation
- S3. Single PTM on the 1$^{st}$ tyrosine site does not yield significant signal peaks in the linker region
- S4. MD simulation on charged interactions between Y3 PTM and MspA bottom arginine residue
- S5. UHPLC-MS results of BCN-C6-TemplateDNA
- S6-S18. UHPLC-MS results of SPPS synthesis and click conjugation of all pentapeptide variants

### Supplementary methods

#### Ultra-high performance liquid chromatography coupled to MS (RP-UHPLC-MS)

Synthesized peptides and peptide-DNA constructs were analyzed by ultra-high performance liquid chromatography (UHPLC) coupled to mass spectrometry (HESI-MS, measuring in negative mode, calibrated with a Thermo Finnigan calibration mixture) using a Q Exactive Focus Agilent 1290 Infinity UHPLC-MS system. Generally, for peptide analysis 10 µL of a 1 mg/mL solution is injected, and for peptide-DNA constructs 10 µL of a 50 µM solution is injected. The UHPLC system is equipped with a diode array detector (DAD G4212A, at 215 nm for peptide detection and 260 nm for DNA detection) and a Dr. Maisch ReproSil Gold 120 C18, 3 µm, 200 x 3 mm column containing a 10 mm guard with a flow rate of 0.4 mL/min. The eluent contained for buffer A: 10 mM triethylammonium acetate in Milli-Q (deionized water, produced with a Milli-Q Integral 3 system; Millipore, Molsheim/France) and for buffer B: acetonitrile (MeCN). A gradient of 5→5→75→75→5→5%, percentage buffer B (0→5→25→28→29→35 min) was applied. Mass spectrometry data is deconvoluted with the use of UniDec software[1].

#### General procedure for Solid-Phase Peptide Synthesis of PSK-like peptides

Peptides were synthesized following Fmoc/*t*Bu Solid-Phase Peptide Synthesis (SPPS) strategy. In general, amino acids were added as follows: the resin was pre-swollen with dichloromethane (DCM). The Fmoc-protecting group was removed using 20% piperidine in *N,N*-dimethylformamide (DMF, 2 x 8 min). The resin was then washed with DMF (3 x 2 min). The next amino acid was treated with 2-(1H-benzotriazole-1-yl)-1,1,3,3-tetramethyluronium hexafluorophosphate (HBTU), 1-hydroxybenzotriazole (HOBt), and *N,N*-diisopropylethylamine (DIPEA) in DMF for 2 min before being added to the resin. The reaction mixture was allowed to couple for 2 h, and reaction completion was monitored by resin staining using ninhydrin (15 g/L, supplemented with 30 mL/L acetic acid in *n*-butanol). Upon reaction completion, the resin was washed with DMF (3 x 2 min), deprotected by 20% piperidine in DMF (2 x 8 min), and washed again with DMF (3 x 2 min). The same steps were repeated until the desired sequence was obtained.

Chain elongation was initiated from Fmoc-Asp(O*t*Bu)-Wang resin 100-200 mesh Novabiochem®, a *p*-alkoxy-benzyl alcohol polymer-bound (copolystyrene-1% DVB) amino acid (loading capacity 0.66 mmol/g). Introduction of new amino acids up until the coupling of Fmoc-Thr(*t*Bu)-OH (14x Asp, 1x Gln, 1x Thr couplings subsequently) was performed in an automatic peptide synthesizer (CS336X Peptide Synthesizer CS BIO Co.). For automated SPPS, a standard protocol was used in which every coupling was reacted for 3h. After automated SPPS, the peptide elongation was continued manually. Beads were split into different SPPS reactors and were either reacted with Fmoc-Tyr(*t*Bu)-OH, Fmoc-Tyr($SO_2$ONp)-OH, or Fmoc-Tyr(PO(OBzl)OH)-OH. After this, Fmoc-Ile-OH was coupled and subsequently beads were again split into multiple SPPS reactors for another round of coupling with either Fmoc-Tyr(*t*Bu)-OH, Fmoc-Tyr($SO_2$ONp)-OH, or Fmoc-Tyr(PO(OBzl)OH)-OH. In the end, azidoacetic acid or azido-$PEG_8$-NHS ester was coupled to the peptides.

After completion of the sequence by SPPS, the peptides were cleaved from the resin by treatment with a cocktail of 90% trifluoroacetic acid (TFA), 2.5% triisopropylsilane (TIS), and 7.5% Milli-Q for 2 h (10 mL TFA cocktail/1 g initial resin) for peptides containing the N-terminal azidoacetic acid moiety. The cleaved peptides were precipitated by dropwise addition of the TFA cocktail-peptide mixture to ice-cold diethyl ether (1:1 ether:hexane, 10x initial cocktail volume) and the cleaved peptide resin was washed once with a small amount of fresh cleavage cocktail. The precipitate was centrifuged for 10 min at 6000 rpm. The supernatant was discarded, and the precipitate was washed with ice-cold diethyl ether and again centrifuged for 10 min at 6000 rpm. This washing step was repeated once more. The resulting precipitate was then dried in a light stream of $N_2$, redissolved in MeCN:Milli-Q (1:1), snap-frozen with liquid $N_2$ and then lyophilized (Labconco FreeZone lyophilizer, 2.5 L, -84 °C, connected to a $35_i$ xDS Edwards Oil-Free Dry Scroll Pump). The lyophilized peptides were treated with 2M $NH_4OAc$ to remove neopentyl from the sulfated tyrosine residues at 45 °C for 40 h. Obtained deprotected peptides were purified by (semi)preparative reverse phase HPLC (Agilent 1260 Preparative HPLC with a DAD G7115A and MSD). Peptides with the azidoacetic acid N-terminus were purified by using a preparative Grace Alltima column (C18, 22 x 250 mm, 5-Micron) with a flow rate of 20 mL/min. The eluent for purification contained for buffer A: 10 mM triethylammonium acetate in Milli-Q, and for buffer B: MeCN. A gradient of 5→10→20→95→95→5→5%, percentage buffer B (0→5→13→14→18→19→20 min) was used. The obtained purified peptides were then lyophilized. Peptides with the azido-$PEG_8$-linker were purified by using a semi-preparative Zorbax Eclipse column (XDB-C18, 9.4 x 250 mm, 5-Micron) with a flow rate of 10 mL/min. The eluent for purification contained for buffer A: 10 mM triethylammonium acetate in Milli-Q, and for buffer B: MeCN. A gradient of 5→5→60→60→5→5%, percentage buffer B (0→5→25→30→35→40 min) was used. The obtained purified peptides were then lyophilized.

**COMSOL simulation on peptide translocation**

Numerical simulations of MspA-helicase-peptide system was implemented in COMSOL Multiphysics 5.4 with a two-dimensional axial symmetrical domain. The simulation included the fluid domain, the lipid membrane, MspA nanopore, helicase, and DNA-linker-peptide strand. The relative permittivity was set to 80, 5 [2], and 3.23 [3] for water, lipid bilayer, and protein (MspA, helicase, and DNA-peptide). The dimensions of M2-MspA nanopore (PDB: 1UUN) were described previously[4]. The charged residues in the inner wall of the nanopore protein were marked at positions of 63 (-), 118 (+), and 134 (+). The geometry of helicase was approximated with a simple ellipsoid with the axial lengths of 5.6 nm, 5.6 nm, and 7.4 nm, based on its outer contour (PDB: 2P6U). It carries -6e (elementary charge) in buffer with pH = 8.0, according to the amino acid sequence. The DNA-linker-peptide strand was approximated with cylindrical columns, with corresponding thickness, located at the central axis of symmetry of the system. The single-strand DNA (ssDNA) carried -1e/nucleotide, the linker was neutral in charge, and the D15 tail carries -1e/amino acid. The charge of tyrosine residues was set at 0, -1e, and -2e, for unmodified, sulfated, and phosphorylated states. The ion distribution and movement in an electrolyte was governed by the Nernst−Planck equation, the electric potential distribution was described by the Poisson equation, and the fluid flow was determined by the Navier−Stokes equations. The *Transport of Diluted Species* module (Nernst−Planck equation), the *Electrostatics* module (Poisson equation), and the *Laminar Flow* module (Navier−Stokes equations) were incorporated and fully coupled in the simulation. The electrolyte was 400 mM KCl with the mobilities of $K^+$ and $Cl^-$ at $7.0 \times 10^{-8}$ and $7.2 \times 10^{-8}$ $m^2\ V^{-1}\ s^{-1}$ respectively[5]. The diffusion coefficients were then determined through the Einstein relation. During the simulation, the DNA-linker-peptide strand was moved upwards with a step size of 0.335 nm, approximating half nucleotide ssDNA step of the helicase[6]. At each step, the steady solutions of corresponding physics fields were found with the bias voltage of 180 mV, in agreement with that used in our experiments. To get the ionic current, the ion flux was integrated on the cross-sectional area located at narrowest restriction of the nanopore.

#### Brief overview of the HMM solver with contamination tolerance

To generate consensus reads from the training set and to align the reads in the test set to the different consensuses, we used a modified hidden Markov model (HMM) solver and a corresponding Baum-Welch expectation-maximization algorithm. Briefly, we sought to accommodate a non-Markovian feature of the nanopore state sequence: the inclusion of spurious "contaminant" states. These contaminant states appear randomly during measurement and violate the Markov property that state transition probabilities depend only on the present state, since the transitions out of a contaminant state are determined by the identity of the most recent non-contaminant state. We therefore derived a maximum *a posteriori* (MAP) algorithm , following the logic of Bahl's previous work[7], while using this "contaminated Markov" property.

In the conventional MAP algorithm, the probability$\alpha_{ti}$, that an observation $X_t$ corresponds to state $i$ given all of the observations $X_1, \dots, X_t$ up to the present, is given by a ``forwards" recursion relation,

$$\alpha_{ti} = \sum_{j} S_{ti} \cdot T_{ji} \cdot \alpha_{t-1,j}$$

where $T_{ji}$ is the probability of transitioning from state $j$ to state $i$ and $S_{ti}$ is the likelihood of observation $X_t$ given the observation distribution associated with state $i$. There is also a similar "backwards" recursion relation for $\beta_{ti}$, the probability that $X_t$ corresponds to state $i$ given the observations $X_{t+1}, \dots, X_n$ , where $X_n$ is the final observation in the sequence. The final likelihood of observation $t$ originating from state $i$ is given by the product $\alpha_{ti}\beta_{ti}$.

In our modified MAP algorithm, we allow probability to carry forward from observations further back than $t - 1$, but subject to the further conditional probability that all the interceding states are identified as contaminants. This results a recursive expression

$$\alpha_{ti} = \sum_{l=1}^{l_{max}} \sum_{j} S_{ti} \cdot T_{ji} \cdot \alpha_{t-l,j} P_c^{l-1}$$

where $P_c$ is the prior probability that any given level is a contaminant. To assign state probabilities, we use this expression together with a corresponding expression for $\beta_{ti}$, as well as for the probability that an observation should be assigned no state and should instead by identified as a contaminant. An implementation of this algorithm is available in the commented MATLAB code titled *fbeMAP.m*. A complete derivation and discussion will be provided in an upcoming publication.

#### Molecular dynamics simulation

For the simulation of the interaction between the nine variants of the PSK-based plant peptide hormones, the following protocol was used in the YASARA software (ref):

1) Construction of the phosphorylated DNA-peptide conjugate according to the molecular structure depicted in Extended Figure 1, up to the oxygen atom of the linker that connects the peptide sequence to the DNA sequence.
2) The mutated MspA nanopore was built using the corresponding coordinates obtained from the Protein Data Bank (PDB-code: 1UUN). The following mutations were

implemented: D134R, E139K, D118R, D90N, D91N, D93N. Then, the nanopore was generated using the oligomerize command.

3) Using the md_membrane macro embedded in the YASARA software package, the nanopore was placed within a biological membrane of which the polar headgroups are identical to those used in the nanopore setup.
4) The peptide was positioned at the middle of the pore. Fix the oxygen atom just above the bottom of the pore. Also fix the nanopore and the membrane. This reduces the simulation time and allows focussing on the effect of the PTMs on the dynamics and interactions of the peptide with the nanopore/membrane.
5) Add a simulation cell around the entire scene.
6) Neutralize the cell at pH 8.0, in the presence of 400 mM KCl (3%) and 10 mM $MgCl_2$ (0.1%), water density: 0.997 g/L.
7) From the same basic structure generated at point 6, all nine variants were generated by swapping atoms (P for S, including automatic adjustment of bond-order, angles and lengths), or deleting the PTM from the tyrosine oxygen atom of the side chain.
8) Each mutant was subjected to a md_run macro for 100 ps.
9) The final structure of the MD simulation of step 8 was subjected to energy minimization to find a local minimum for the peptide-nanopore interaction.

**SI Table 1**. Sequences of DNA and DNA-peptide conjugates in this study.

| Name | Sequence information |
|---|---|
| Template DNA | [5'BCN]TTACTGAAGTCTCACGTGCCTGGTATATTAGCGTCCACTCTCACTATCGGAT-TCTACATCGGTCGTAGCC |
| Complement DNA | CCGATGTAGAATCCGATAGTGAGAGTTTTTTTTTTTTTTTTTTTT [3'cholesterol] |
| JS445 | TemplateDNA-BCN-triazole-YIYTQ-DDDDD-DDDDD-DDDDD |
| JS446 | TemplateDNA-BCN-triazole-Y[$SO_3$]IYTQ-DDDDD-DDDDD-DDDDD |
| JS447 | TemplateDNA-BCN-triazole-YIY[$SO_3$]TQ-DDDDD-DDDDD-DDDDD |
| JS448 | TemplateDNA-BCN-triazole-Y[$SO_3$]IY[$SO_3$]TQ-DDDDD-DDDDD-DDDDD |
| JS449 | TemplateDNA-BCN-triazole-Y[$SO_3$]IY[$PO_3$]TQ-DDDDD-DDDDD-DDDDD |
| JS450 | TemplateDNA-BCN-triazole-Y[$PO_3$]IY[$SO_3$]TQ-DDDDD-DDDDD-DDDDD |
| JS451 | TemplateDNA-BCN-triazole-Y[$PO_3$]IYTQ-DDDDD-DDDDD-DDDDD |
| JS452 | TemplateDNA-BCN-triazole-YIY[$PO_3$]TQ-DDDDD-DDDDD-DDDDD |
| JS453 | TemplateDNA-BCN-triazole-Y[$PO_3$]IY[$PO_3$]TQ-DDDDD-DDDDD-DDDDD |
| JS527 | TemplateDNA-BCN-triazole-PEG8-YIYTQ-DDDDD-DDDDD-DDDDD |
| JS528 | TemplateDNA-BCN-triazole-PEG8-YIY[$SO_3$]TQ-DDDDD-DDDDD-DDDDD |
| JS529 | TemplateDNA-BCN-triazole-PEG8-Y[$SO_3$]IYTQ-DDDDD-DDDDD-DDDDD |
| JS530 | TemplateDNA-BCN-triazole-PEG8-Y[$SO_3$]IY[$SO_3$]TQ-DDDDD-DDDDD-DDDDD |

### Supplementary figures

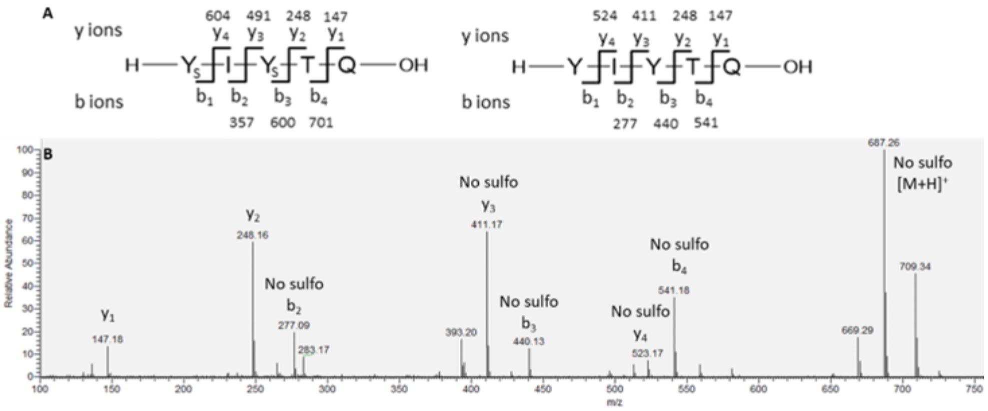


**Supplementary Figure S1.** CID results on sulfation under mass spec. (A) Calculated mass spectrometry fragments of PSK and non-sulfated PSK. (B) Fragmentation of PSK by collision-induced dissociation (CID) where a complete y and b-type ion series was observed while $SO_3$ was detectably lost from the fragmented ions or the precursor ion.

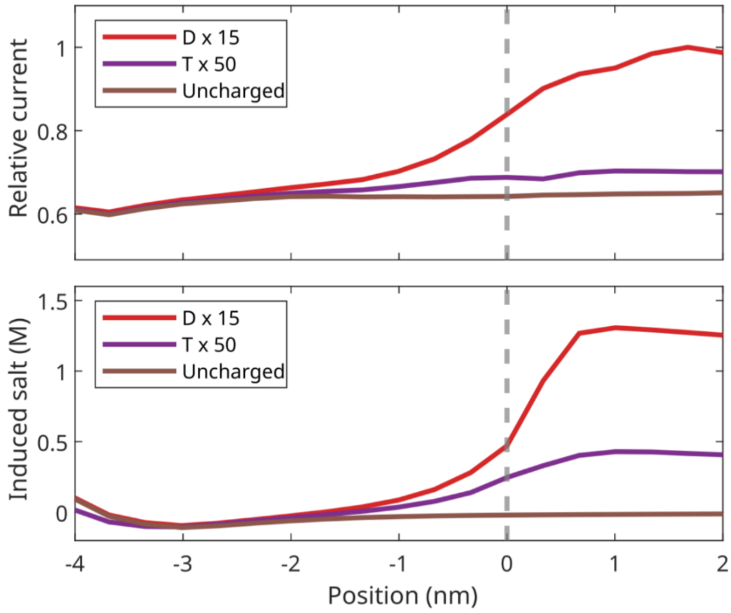


**Supplementary Figure S2.** COMSOL simulation on charge-induced salt concentration and ionic current elevation. Poly-T ssDNA tail only shows minor signal elevation in the simulation. The density of the charges is critical for the salt modulation effect.

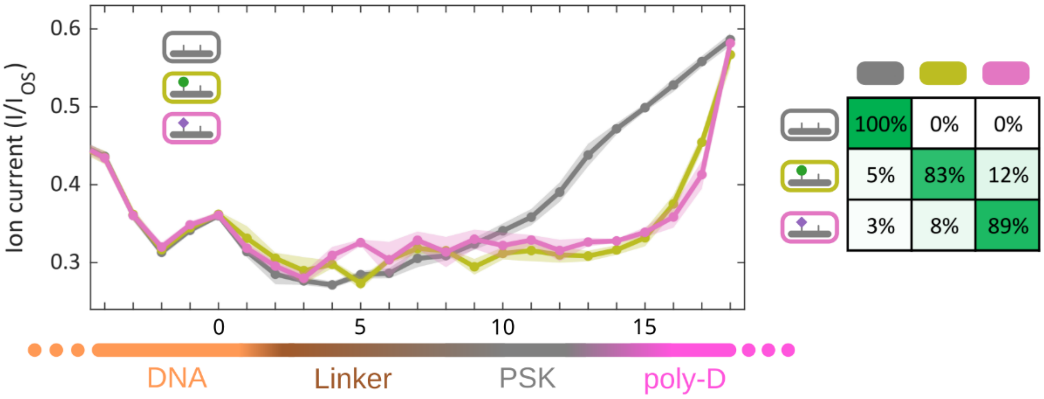


**Supplementary Figure S3.** Single PTM on the 1st tyrosine site does not yield significant signal peaks in the linker region. The influence of the PTM is reflected by the delay of the signal ramp. The variant calling accuracies are still high because of the overall higher signals in phosphorylated samples.

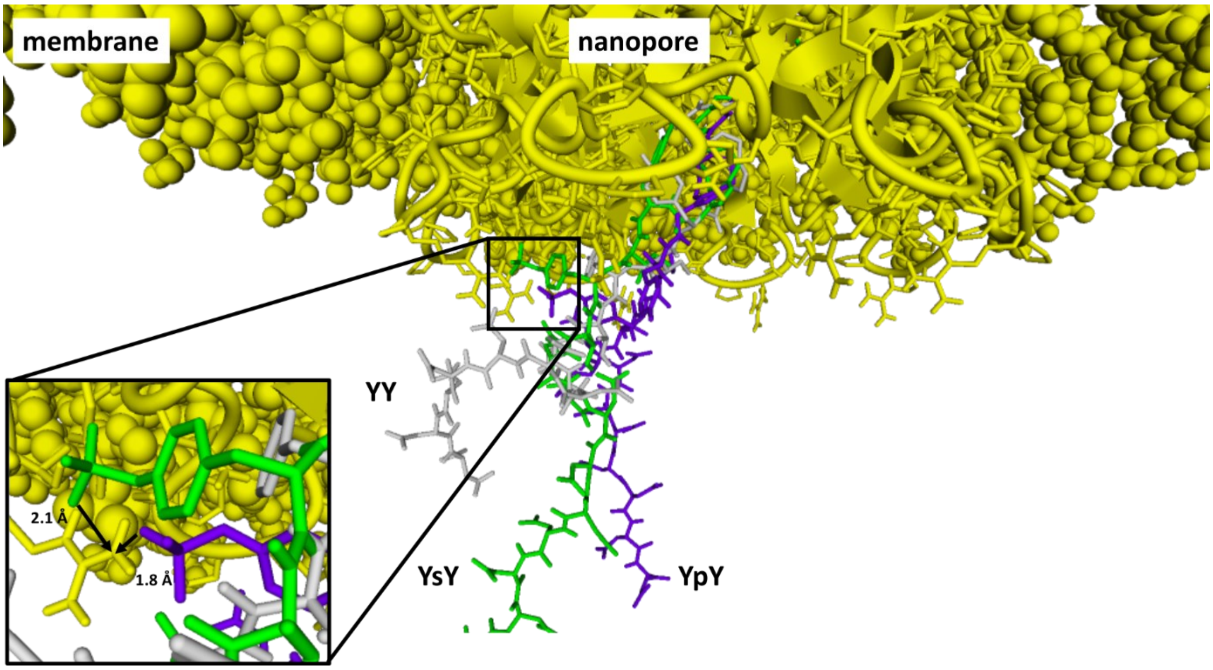


**Supplementary Figure S4.** MD simulation on charged interactions between Y3 PTM and MspA bottom arginine residue. The PSK sequence containing unmodified Tyr residues (grey) adopts a more compact structure, whereas the sequences with a sulfotyrosine (green) or phosphotyrosine (purple) on the 2nd tyrosine residue interact with the arginine residues at the mouth of the MspA pore (distances are given between oxygen atom on sY or pY and most proximal NH2-group of the arginine side-chain).

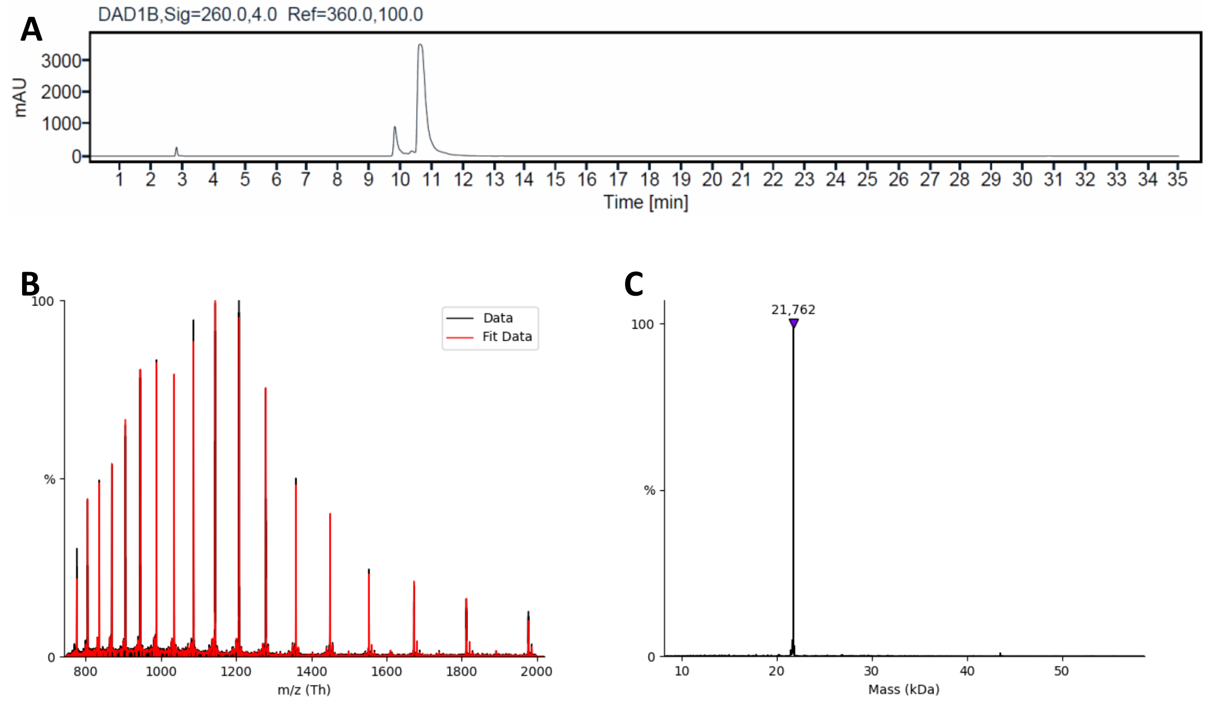


**Supplementary Figure S5**. (A) UHPLC trace of BCN-C6-TemplateDNA. The tR of the DNA is 10.6 min, and is >89% pure; the peak at $t_R$ <10 min belongs to DNA that does not contain a BCN handle. (B) Multiply charged ion series of MS spectrum in negative mode. (C) Deconvoluted HESI mass spectrum of BCN-C6-TemplateDNA. Expected mass 21763, observed 21762.

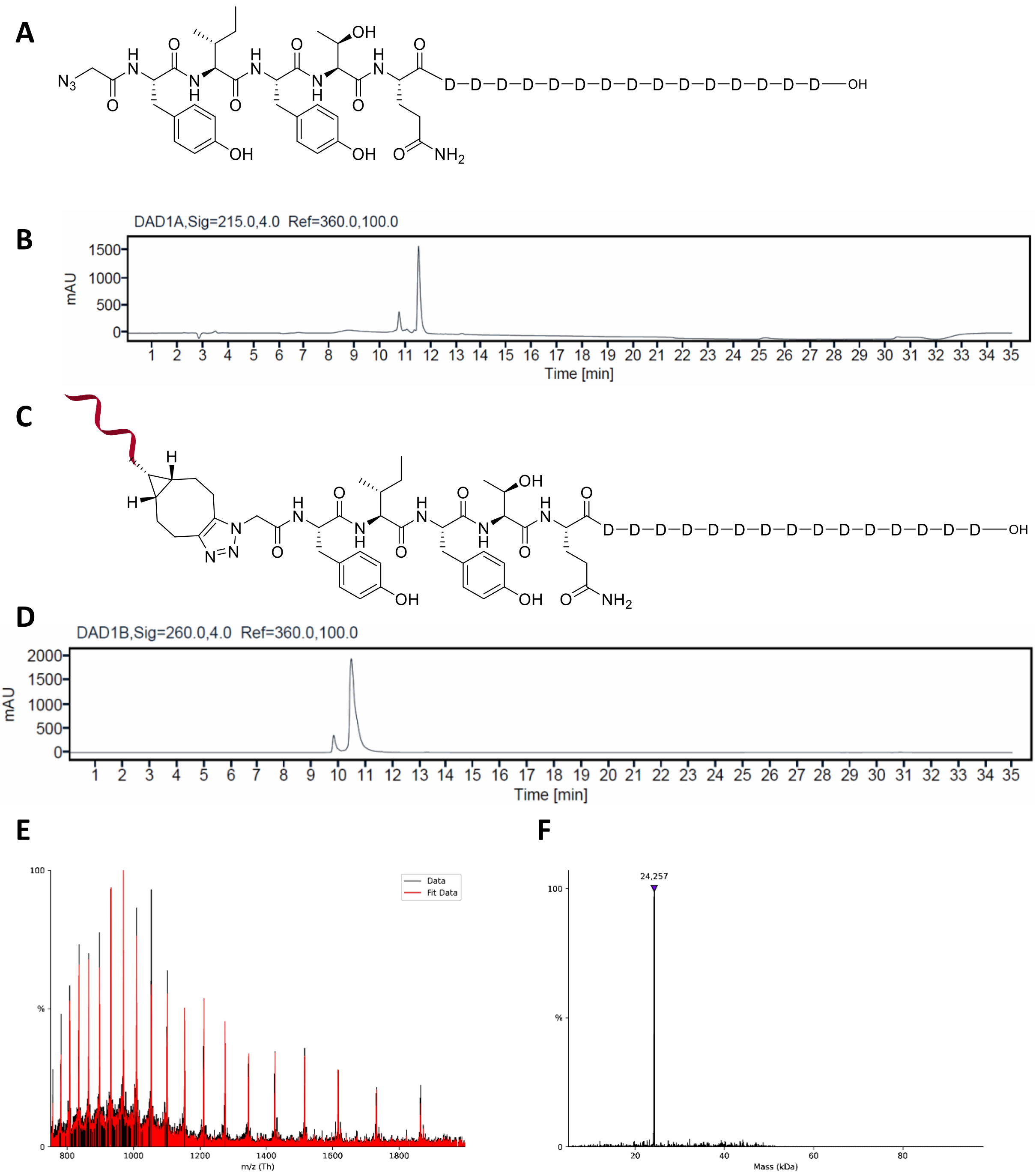


**Supplementary Figure S6**. (A) Chemical structure of azido-methyl-Tyr-Ile-Tyr-Thr-Gln-(Asp)$_{15}$-OH. (B) UHPLC trace, the $t_R$ of the product is 11.5 min. HRMS (ESI): $m/z$ = [M-2H]$^{2-}$ calc for $C_{95}H_{120}N_{24}O_{56}$ 1246.3645, found 1246.3671; $m/z$ = [M-3H]$^{3-}$ calc for $C_{95}H_{119}N_{24}O_{56}$ 830.5739, found 830.5768. (C) Chemical structure of Template DNA-methyl-Tyr-Ile-Tyr-Thr-Gln-(Asp)$_{15}$-OH. (D) UHPLC trace, the $t_R$ of the product is 10.5 min. (E) Multiply charged ion series of MS spectrum in negative mode. (F) Deconvoluted HESI mass spectrum. Expected mass 24258, observed 24257.

**Template DNA-methyl-Tyr(SO3H)-Ile-Tyr-Thr-Gln-(Asp)15-OH**

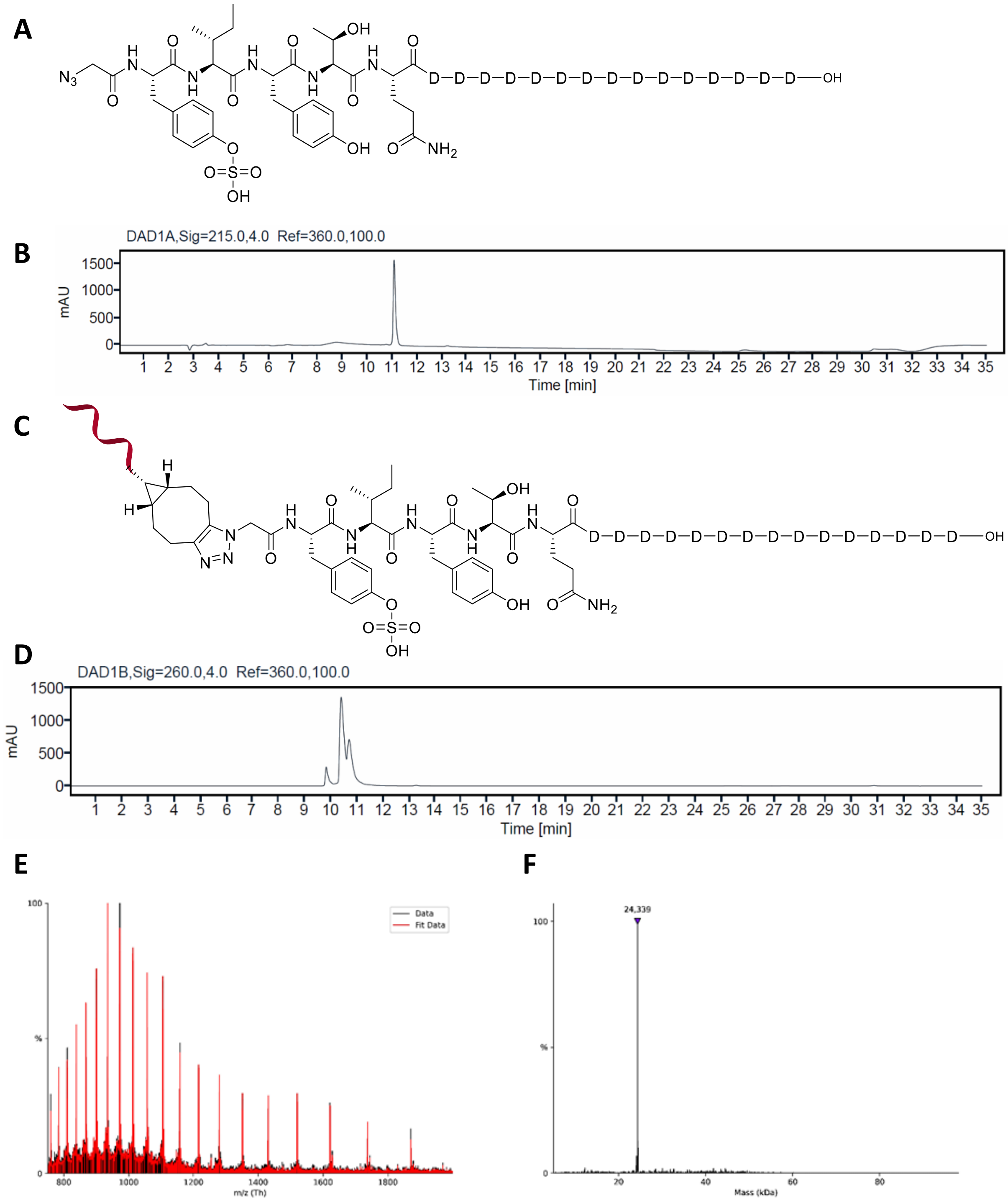


**Supplementary Figure S7.** (A) Chemical structure of azido-methyl-Tyr($SO_3$H)-Ile-Tyr-Thr-Gln-(Asp)$_{15}$-OH. (B) UHPLC trace, the $t_R$ of the product is 11.1 min. HRMS (ESI): *m/z* = [M-2H]$^{2-}$ calc for $C_{95}H_{120}N_{24}O_{59}S$ 1286.3430, found 1286.3462; *m/z* = [M-3H]$^{3-}$ calc for $C_{95}H_{119}N_{24}O_{59}S$ 857.2262, found 857.2293. (C) Chemical structure of Template DNA-methyl-Tyr($SO_3$H)-Ile-Tyr-Thr-Gln-(Asp)$_{15}$-OH. (D) UHPLC trace, the $t_R$ of the product is 10.4 min. (E) Multiply charged ion series of MS spectrum in negative mode for the peak at $t_R$ at 10.4 min. (F) Deconvoluted HESI mass spectrum. Expected mass 24339, observed 24339.

**Template DNA-methyl-Tyr-Ile-Tyr($SO_3H$)-Thr-Gln-(Asp)$_{15}$-OH**

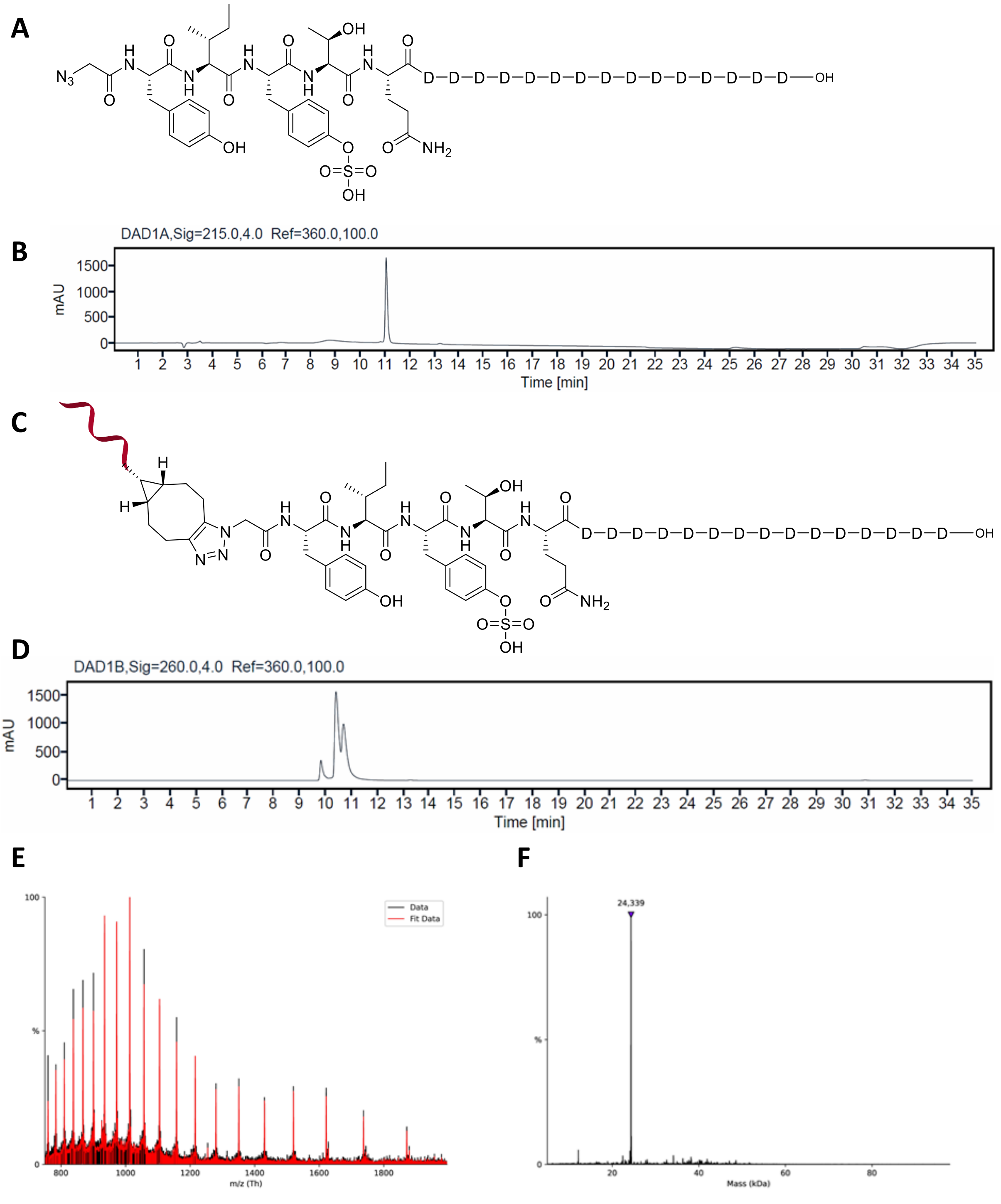

**Supplementary Figure S8.** (A) Chemical structure of azido-methyl-Tyr-Ile-Tyr($SO_3H$)-Thr-Gln-(Asp)$_{15}$-OH. (B) UHPLC trace, the $t_R$ of the product is 11.0 min. HRMS (ESI): *m/z* = $[M-2H]^{2-}$ calc for $C_{95}H_{120}N_{24}O_{59}S$ 1286.3430, found 1286.3465; *m/z* = $[M-3H]^{3-}$ calc for $C_{95}H_{119}N_{24}O_{59}S$ 857.2262, found 857.2298. (C) Chemical structure of Template DNA-methyl-Tyr-Ile-Tyr($SO_3H$)-Thr-Gln-(Asp)$_{15}$-OH. (D) UHPLC trace, the $t_R$ of the product is 10.4 min. (E) Multiply charged ion series of MS spectrum in negative mode for the peak at $t_R$ at 10.4 min. (F) Deconvoluted HESI mass spectrum. Expected mass 24339, observed 24339.

**Template DNA-methyl-Tyr(SO$_3$H)-Ile-Tyr(SO$_3$H)-Thr-Gln-(Asp)$_{15}$-OH**

**A**

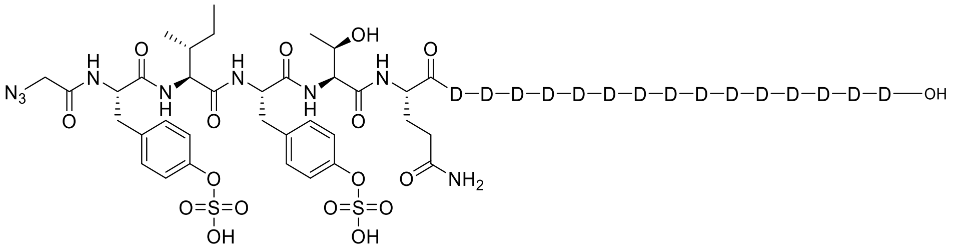


**B**

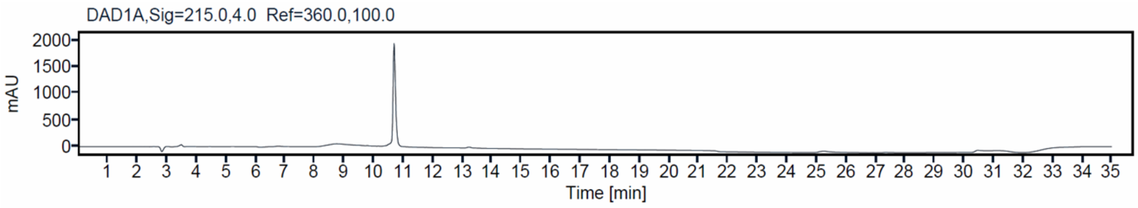


**C**

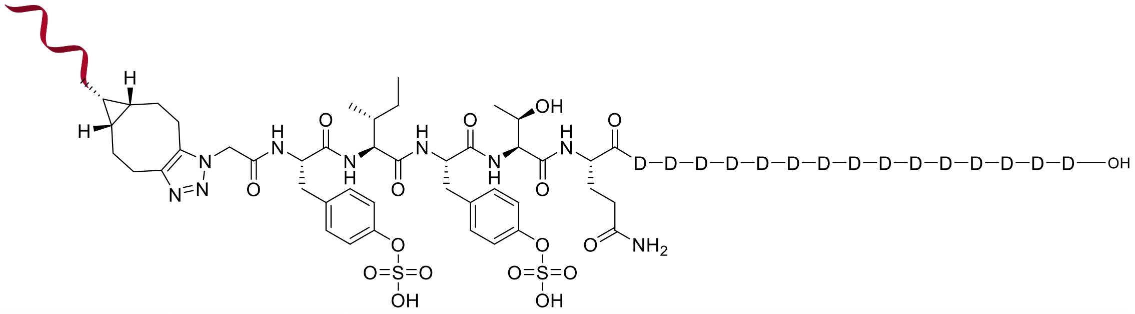


**D**

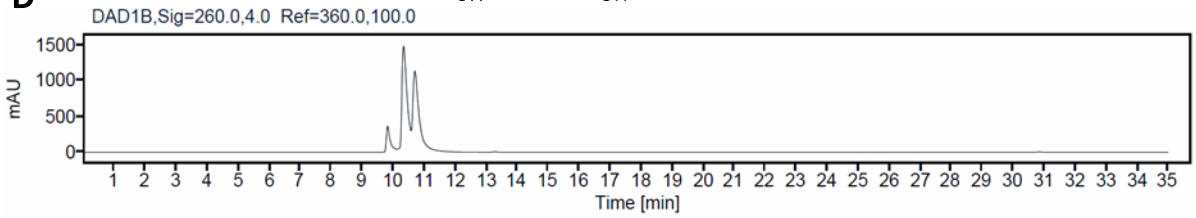


**E**

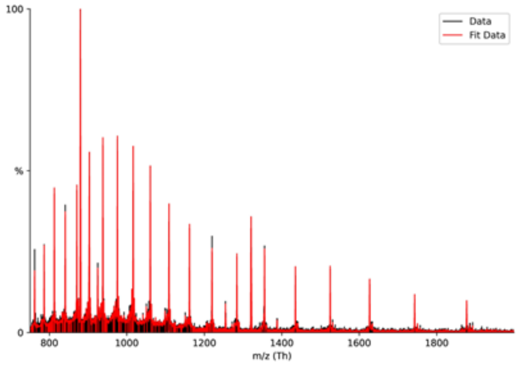


**F**

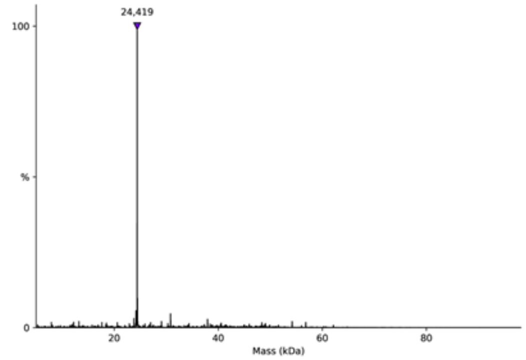


**Supplementary Figure S9.** (A) Chemical structure of azido-methyl-Tyr(SO$_3$H)-Ile-Tyr(SO$_3$H)-Thr-Gln-(Asp)$_{15}$-OH. (B) UHPLC trace, the $t_R$ of the product is 10.7 min. HRMS (ESI): *m/z* = [M-2H]$^{2-}$ calc for $C_{95}H_{120}N_{24}O_{62}S_2$ 1326.3214, found 1326.3246; *m/z* = [M-3H]$^{3-}$ calc for $C_{95}H_{119}N_{24}O_{62}S_2$ 883.8785, found 883.8822. (C) Chemical structure of Template DNA-methyl-Tyr(SO$_3$H)-Ile-Tyr(SO$_3$H)-Thr-Gln-(Asp)$_{15}$-OH. (D) UHPLC trace, the $t_R$ of the product is 10.4 min. (E) Multiply charged ion series of MS spectrum in negative mode for the peaks at $t_R$ 10.4 and 10.7 min. (F) Deconvoluted HESI mass spectrum. Expected mass 24419, observed 24419.

**Template DNA-methyl-Tyr($PO_3H_2$)-Ile-Tyr-Thr-Gln-(Asp)$_{15}$-OH**

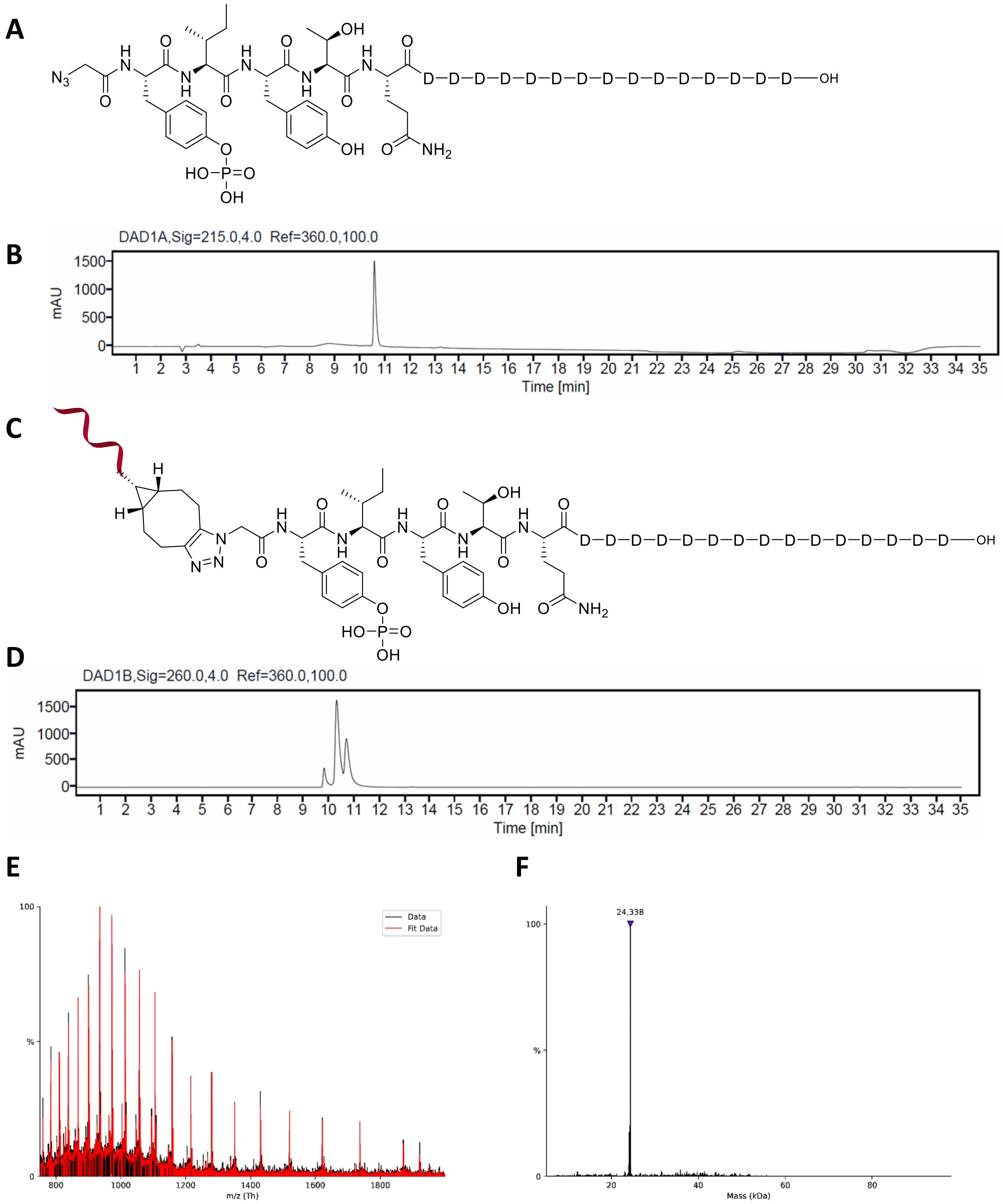


**Supplementary Figure S10.** (A) Chemical structure of azido-methyl-Tyr($PO_3H_2$)-Ile-Tyr-Thr-Gln-(Asp)$_{15}$-OH. (B) UHPLC trace, the $t_R$ of the product is 10.6 min. HRMS (ESI): *m/z* = $[M-2H]^{2-}$ calc for $C_{95}H_{121}N_{24}O_{59}P$ 1286.3477, found 1286.3516; *m/z* = $[M-3H]^{3-}$ calc for $C_{95}H_{120}N_{24}O_{59}P$ 857.2294, found 857.2333. (C) Chemical structure of Template DNA-methyl-Tyr($PO_3H_2$)-Ile-Tyr-Thr-Gln-(Asp)$_{15}$-OH. (D) UHPLC trace, the $t_R$ of the product is 10.3 min. (E) Multiply charged ion series of MS spectrum in negative mode for the peaks at $t_R$ 10.3 and 10.7 min. (F) Deconvoluted HESI mass spectrum. Expected mass 24339, observed 24338.

**Template DNA-methyl-Tyr-Ile-Tyr($PO_3H_2$)-Thr-Gln-(Asp)$_{15}$-OH**

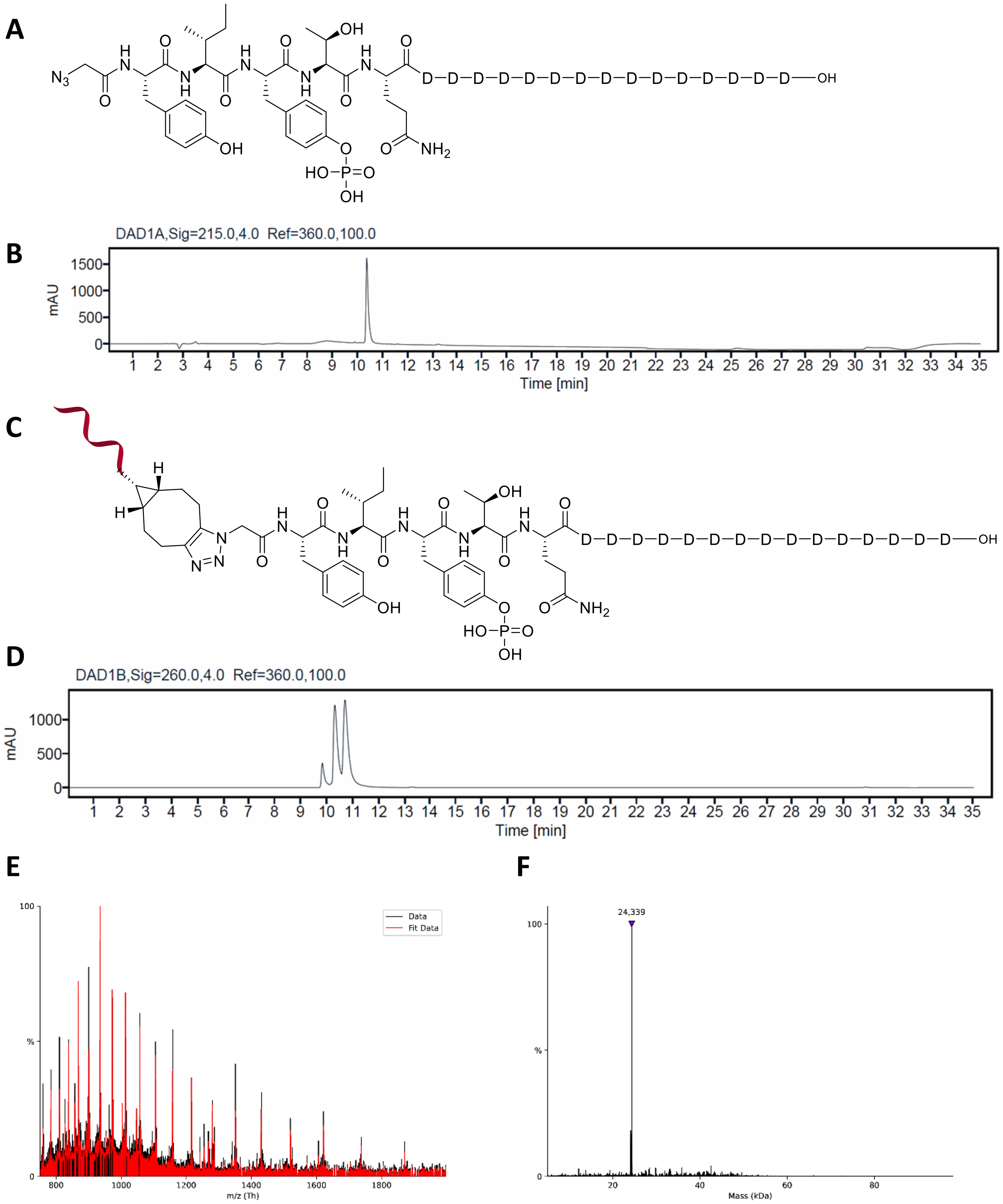


**Supplementary Figure S11.** (A) Chemical structure of azido-methyl-Tyr-Ile-Tyr($PO_3H_2$)-Thr-Gln-(Asp)$_{15}$-OH. (B) UHPLC trace, the $t_R$ of the product is 10.3 min. HRMS (ESI): $m/z$ = $[M-2H]^{2-}$ calc for $C_{95}H_{121}N_{24}O_{59}P$ 1286.3477, found 1286.3517; $m/z$ = $[M-3H]^{3-}$ calc for $C_{95}H_{120}N_{24}O_{59}P$ 857.2294, found 857.2332. (C) Chemical structure of Template DNA-methyl-Tyr-Ile-Tyr($PO_3H_2$)-Thr-Gln-(Asp)$_{15}$-OH. (D) UHPLC trace, the $t_R$ of the product is 10.3 min. (E) Multiply charged ion series of MS spectrum in negative mode for the peaks at $t_R$ 10.3 and 10.7 min. (F) Deconvoluted HESI mass spectrum. Expected mass 24339, observed 24339.

**Template DNA-methyl-Tyr($PO_3H_2$)-Ile-Tyr($PO_3H_2$)-Thr-Gln-(Asp)$_{15}$-OH**

**A**

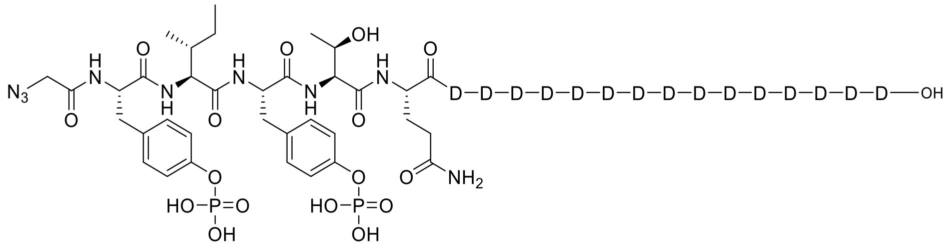


**B**

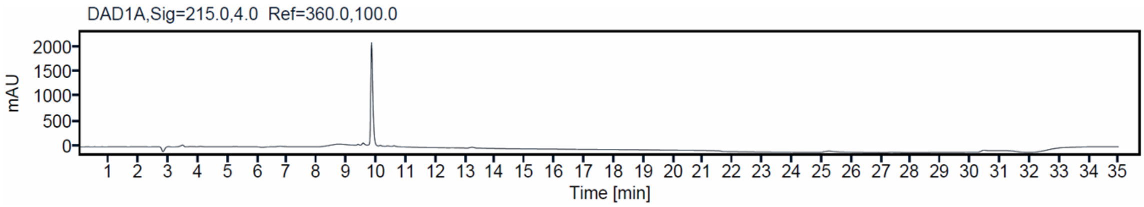


**C**

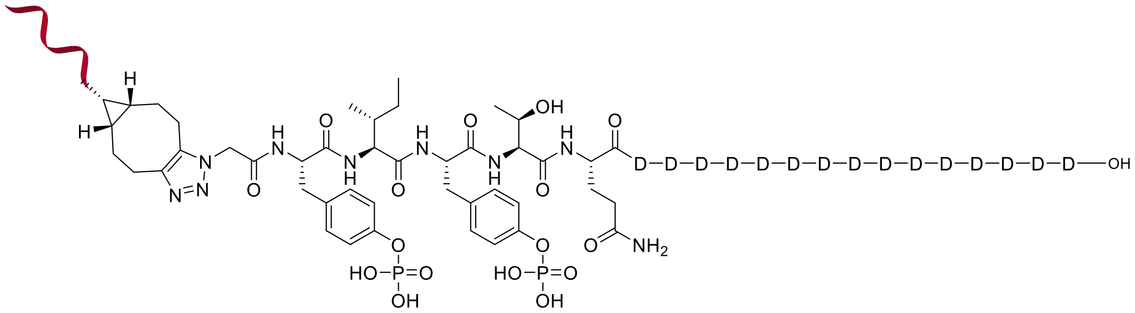


**D**

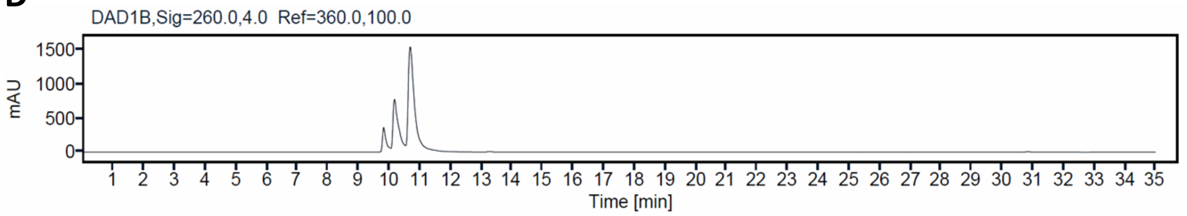


**E**

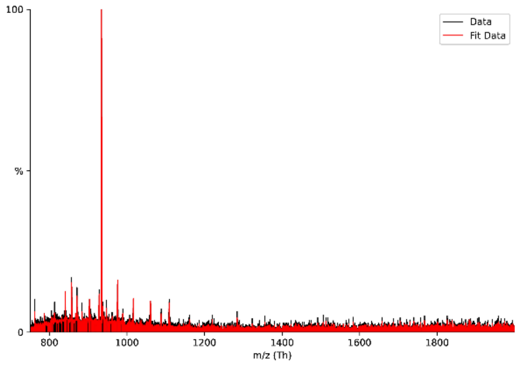


**F**

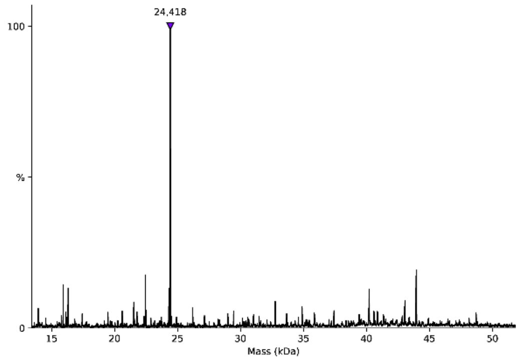


**Supplementary Figure S12.** (A) Chemical structure of azido-methyl-Tyr($PO_3H_2$)-Ile-Tyr($PO_3H_2$)-Thr-Gln-(Asp)$_{15}$-OH. (B) UHPLC trace, the $t_R$ of the product is 9.8 min. HRMS (ESI): *m/z* = $[M-2H]^{2-}$ calc for $C_{95}H_{122}N_{24}O_{62}P_2$ 1326.3309, found 1326.3338; *m/z* = $[M-3H]^{3-}$ calc for $C_{95}H_{121}N_{24}O_{62}P_2$ 883.8848, found 883.8884. (C) Chemical structure of Template DNA-methyl-Tyr($PO_3H_2$)-Ile-Tyr($PO_3H_2$)-Thr-Gln-(Asp)$_{15}$-OH. (D) UHPLC trace, the $t_R$ of the product is 10.2 min. (E) Multiply charged ion series of MS spectrum in negative mode for the peaks at $t_R$ 10.2 and 10.7 min. (F) Deconvoluted HESI mass spectrum. Expected mass 24419, observed 24418.

**Template DNA-methyl-Tyr($SO_3H$)-Ile-Tyr($PO_3H_2$)-Thr-Gln-(Asp)$_{15}$-OH**

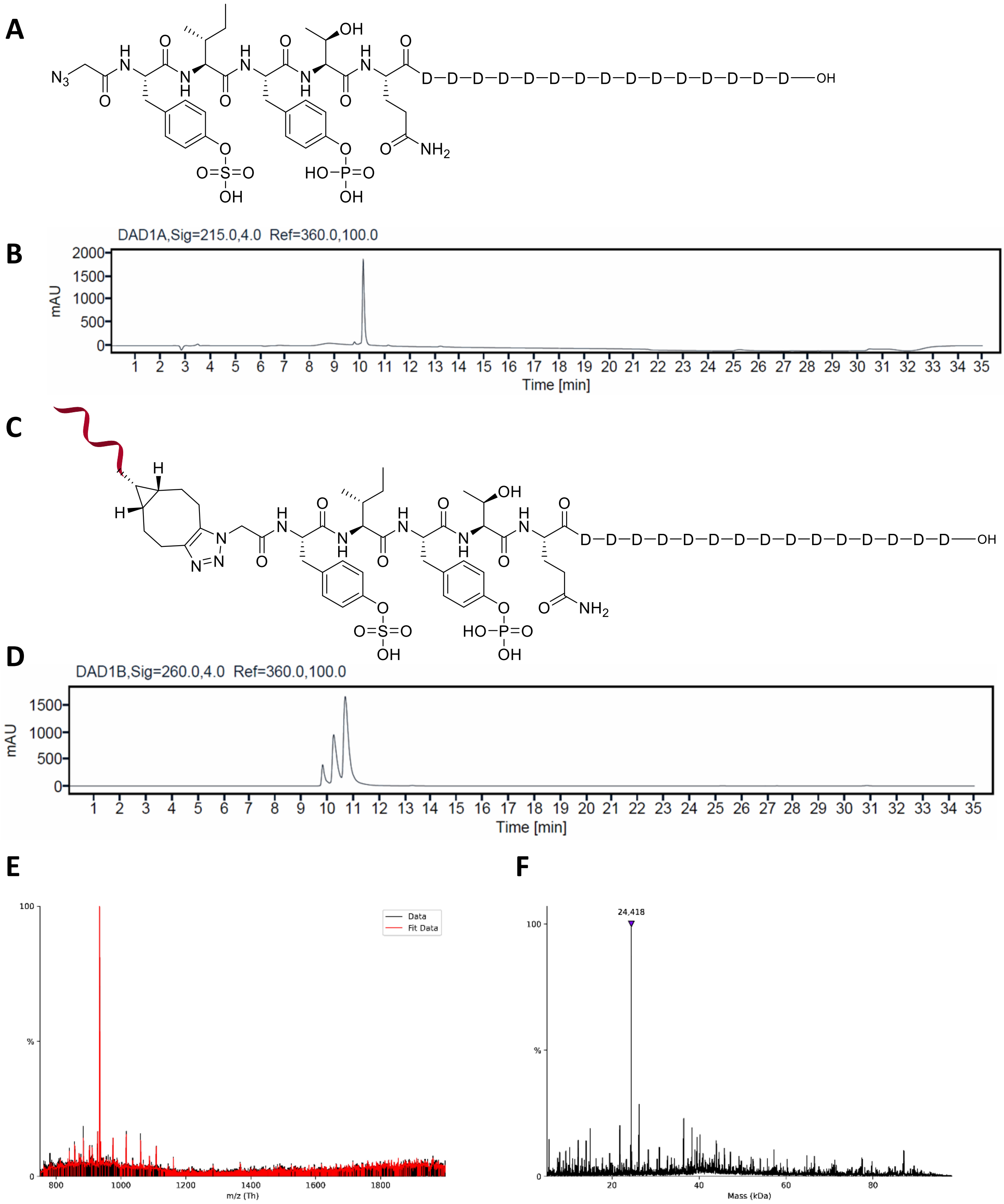


**Supplementary Figure S13.** (A) Chemical structure of azido-methyl-Tyr($SO_3H$)-Ile-Tyr($PO_3H_2$)-Thr-Gln-(Asp)$_{15}$-OH. (B) UHPLC trace, the $t_R$ of the product is 10.1 min. HRMS (ESI): $m/z$ = $[M-2H]^{2-}$ calc for $C_{95}H_{121}N_{24}O_{62}PS$ 1326.3261, found 1326.3297; $m/z$ = $[M-3H]^{3-}$ calc for $C_{95}H_{120}N_{24}O_{62}PS$ 883.8817, found 883.8857. (C) Chemical structure of Template DNA-methyl-Tyr($SO_3H$)-Ile-Tyr($PO_3H_2$)-Thr-Gln-(Asp)$_{15}$-OH. (D) UHPLC trace, the $t_R$ of the product is 10.2 min. (E) Multiply charged ion series of MS spectrum in negative mode for the peaks at $t_R$ 10.2 and 10.7 min. (F) Deconvoluted HESI mass spectrum. Expected mass 24419, observed 24418.

**Template DNA-methyl-Tyr($PO_3H_2$)-Ile-Tyr($SO_3H$)-Thr-Gln-(Asp)$_{15}$-OH**

**A**

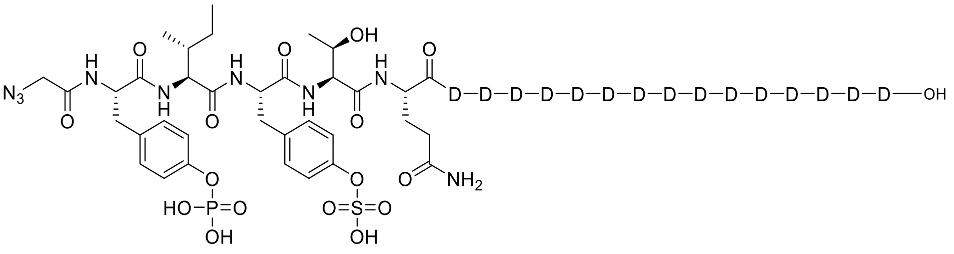

**B**

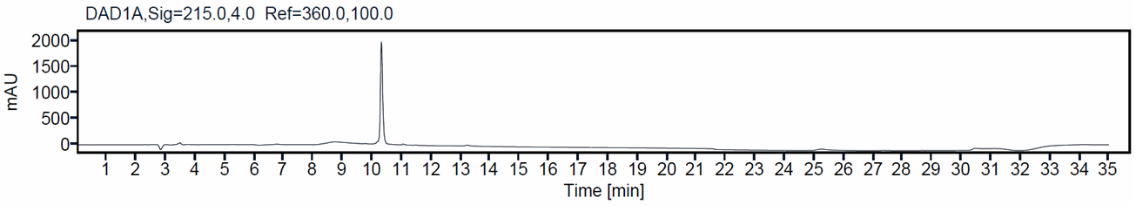


**C**

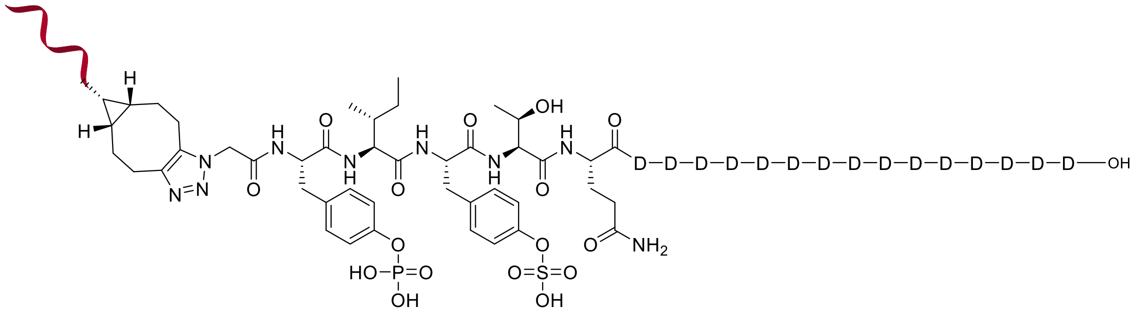

**D**

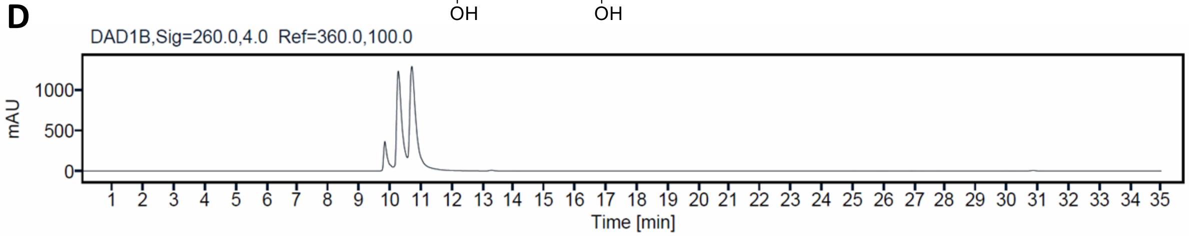


**E**

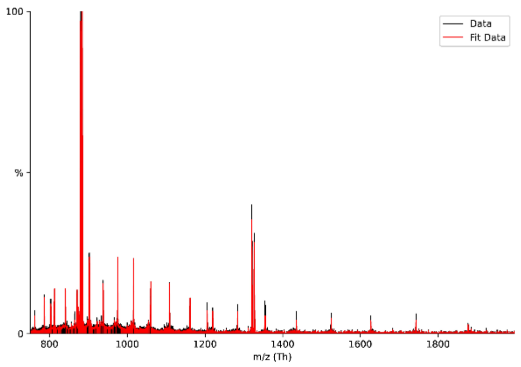


**F**

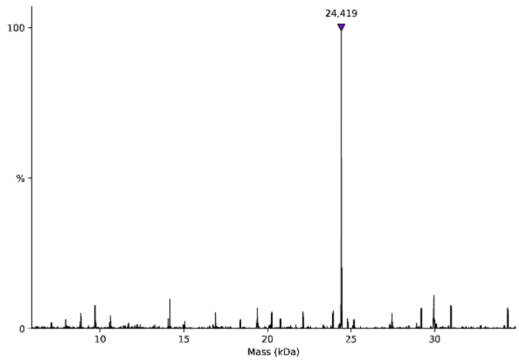


**Supplementary Figure S14.** (A) Chemical structure of azido-methyl-Tyr($PO_3H_2$)-Ile-Tyr($SO_3H$)-Thr-Gln-(Asp)$_{15}$-OH. (B) UHPLC trace, the $t_R$ of the product is 10.3 min. HRMS (ESI): $m/z$ = $[M-2H]^{2-}$ calc for $C_{95}H_{121}N_{24}O_{62}PS$ 1326.3261, found 1326.3295; $m/z$ = $[M-3H]^{3-}$ calc for $C_{95}H_{120}N_{24}O_{62}PS$ 883.8817, found 883.8854. (C) Chemical structure of Template DNA-methyl-Tyr($PO_3H_2$)-Ile-Tyr($SO_3H$)-Thr-Gln-(Asp)$_{15}$-OH. (D) UHPLC trace, the $t_R$ of the product is 10.2 min. (E) Multiply charged ion series of MS spectrum in negative mode for the peaks at $t_R$ 10.2 and 10.7 min. (F) Deconvoluted HESI mass spectrum. Expected mass 24419, observed 24419.

#### Template DNA-PEG$_8$-Tyr-Ile-Tyr-Thr-Gln-(Asp)$_{15}$-OH

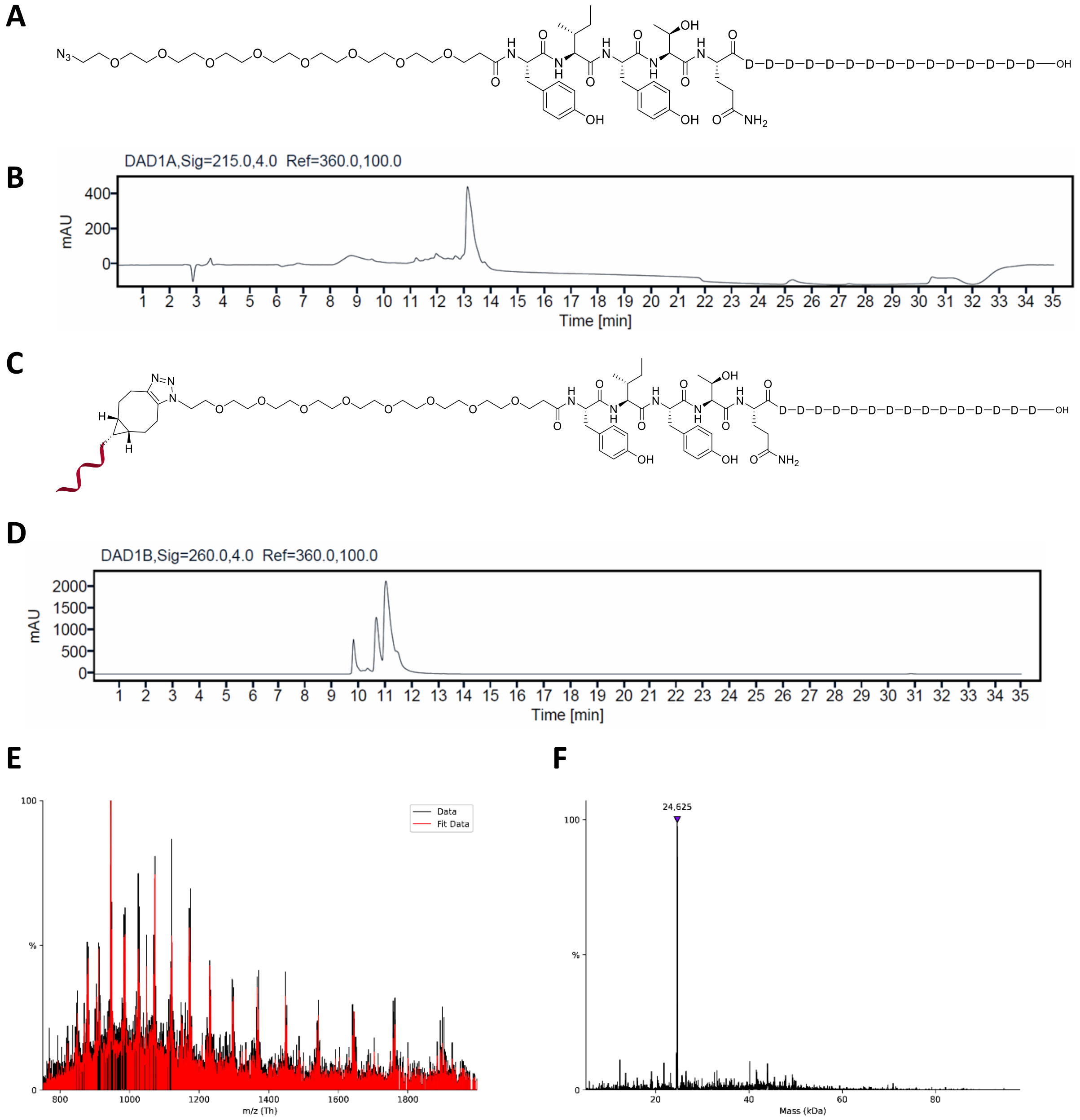


**Supplementary Figure S15.** (A) Chemical structure of azido-PEG$_8$-Tyr-Ile-Tyr-Thr-Gln-(Asp)$_{15}$-OH. (B) UHPLC trace, the $t_R$ of the product is 13.1 min. HRMS (ESI): *m/z* = $[M-2H]^{2-}$ calc for $C_{112}H_{154}N_{24}O_{64}$ 1429.4772, found 1429.4825; *m/z* = $[M-3H]^{3-}$ calc for $C_{112}H_{153}N_{24}O_{64}$ 952.6491, found 952.6526. (C) Chemical structure of Template DNA-PEG$_8$-Tyr-Ile-Tyr-Thr-Gln-(Asp)$_{15}$-OH. (D) UHPLC trace, the $t_R$ of the product is 11.0 min. (E) Multiply charged ion series of MS spectrum in negative mode. (F) Deconvoluted HESI mass spectrum. Expected mass 24625, observed 24625.

#### Template DNA-PEG$_8$-Tyr(SO$_3$H)-Ile-Tyr-Thr-Gln-(Asp)$_{15}$-OH

**A**

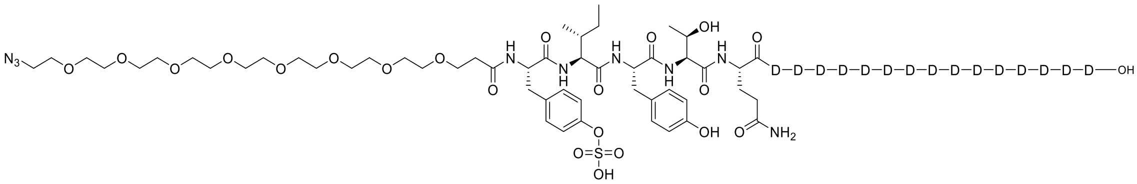

**B**

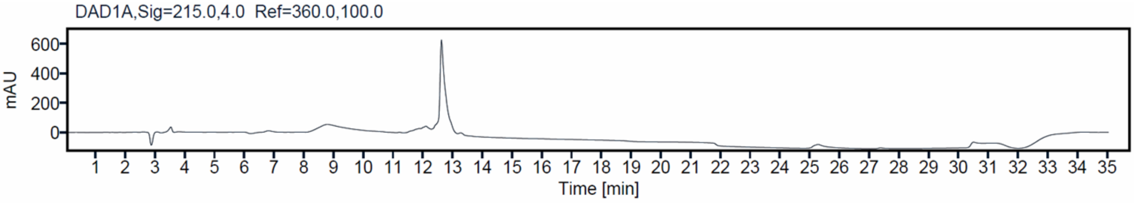


**C**

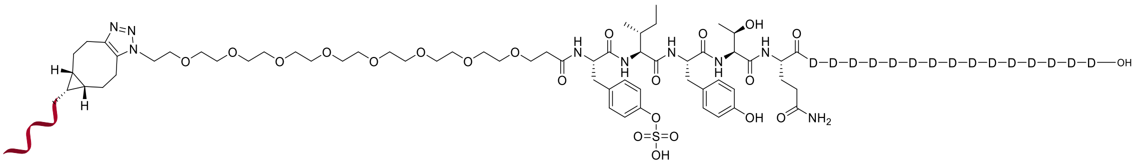

**D**

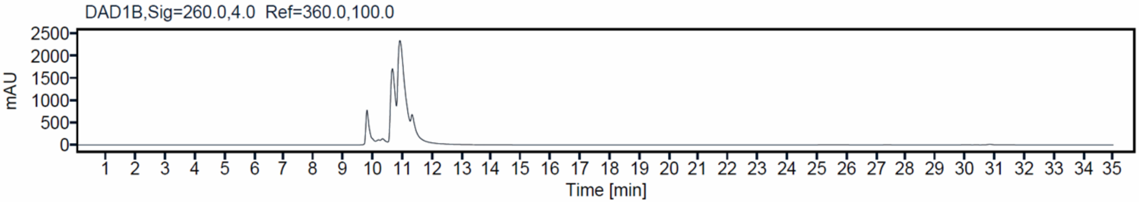


**E**

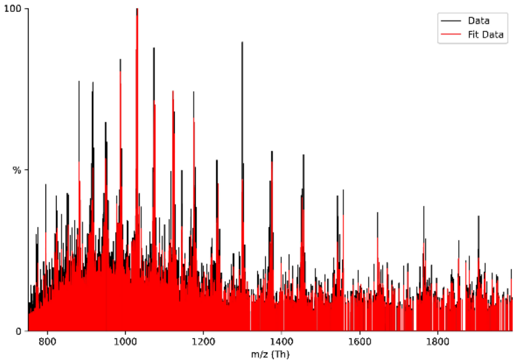


**F**

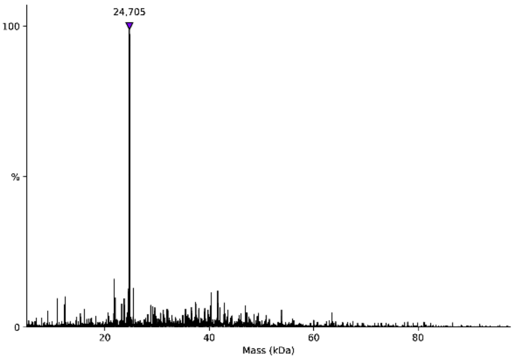


**Supplementary Figure S16.** (A) Chemical structure of azido-PEG$_8$-Tyr(SO$_3$H)-Ile-Tyr-Thr-Gln-(Asp)$_{15}$-OH. (B) UHPLC trace, the $t_R$ of the product is 12.6 min. HRMS (ESI): *m/z* = [M-2H]$^{2-}$ calc for $C_{112}H_{154}N_{24}O_{67}S$ 1469.4556, found 1469.4599; *m/z* = [M-3H]$^{3-}$ calc for $C_{112}H_{153}N_{24}O_{67}S$ 979.3013, found 979.3049. (C) Chemical structure of Template DNA-PEG$_8$-Tyr(SO$_3$H)-Ile-Tyr-Thr-Gln-(Asp)$_{15}$-OH. (D) UHPLC trace, the $t_R$ of the product is 10.9 min. (E) Multiply charged ion series of MS spectrum in negative mode. (F) Deconvoluted HESI mass spectrum. Expected mass 24705, observed 24705.

**Template DNA-PEG$_8$-Tyr-Ile-Tyr(SO$_3$H)-Thr-Gln-(Asp)$_{15}$-OH**

**A**

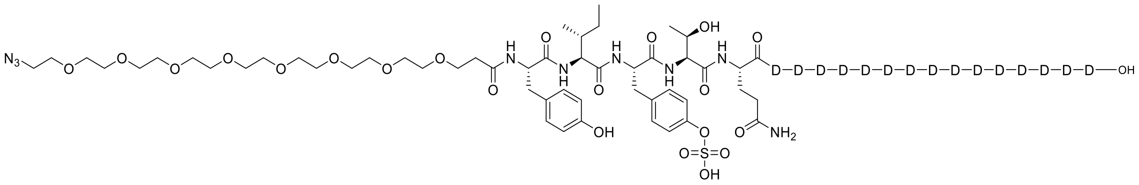

**B**

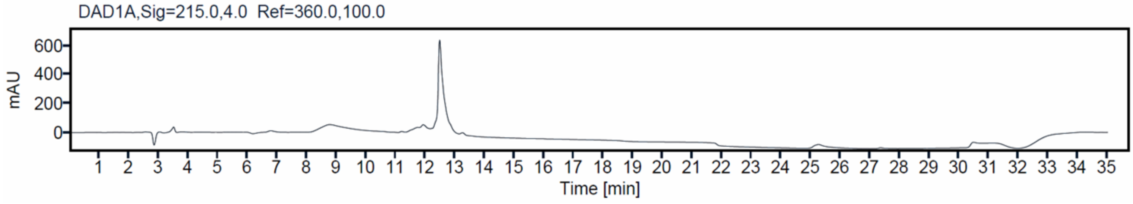


**C**

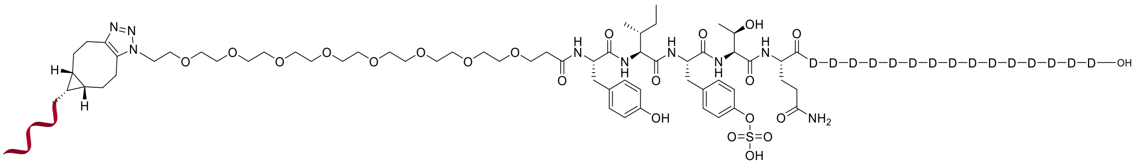

**D**

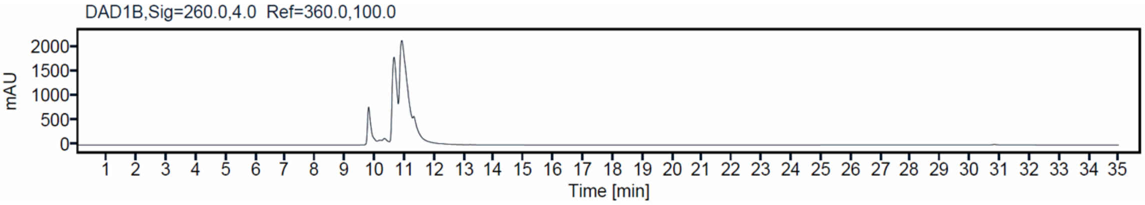


**E**

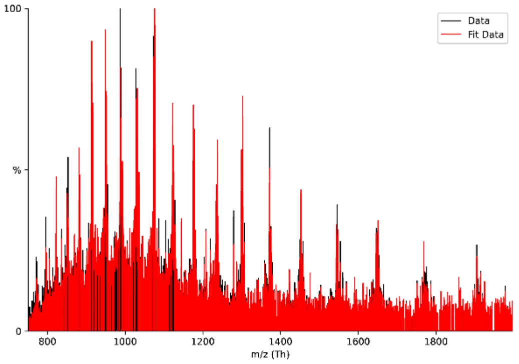


**F**

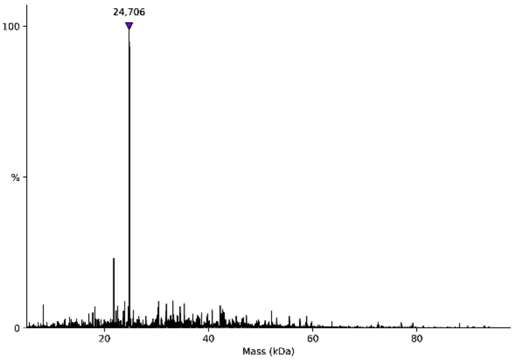


**Supplementary Figure S17.** (A) Chemical structure of azido-PEG$_8$-Tyr-Ile-Tyr(SO$_3$H)-Thr-Gln-(Asp)$_{15}$-OH. (B) UHPLC trace, the $t_R$ of the product is 12.5 min. HRMS (ESI): *m/z* = $[M-2H]^{2-}$ calc for $C_{112}H_{154}N_{24}O_{67}S$ 1469.4556, found 1469.4597; *m/z* = $[M-3H]^{3-}$ calc for $C_{112}H_{153}N_{24}O_{67}S$ 979.3013, found 979.3047. (C) Chemical structure of Template DNA-PEG$_8$-Tyr-Ile-Tyr(SO$_3$H)-Thr-Gln-(Asp)$_{15}$-OH. (D) UHPLC trace, the $t_R$ of the product is 10.9 min. (E) Multiply charged ion series of MS spectrum in negative mode. (F) Deconvoluted HESI mass spectrum. Expected mass 24705, observed 24706.

**Template DNA-PEG$_8$-Tyr(SO$_3$H)-Ile-Tyr(SO$_3$H)-Thr-Gln-(Asp)$_{15}$-OH**

**A**

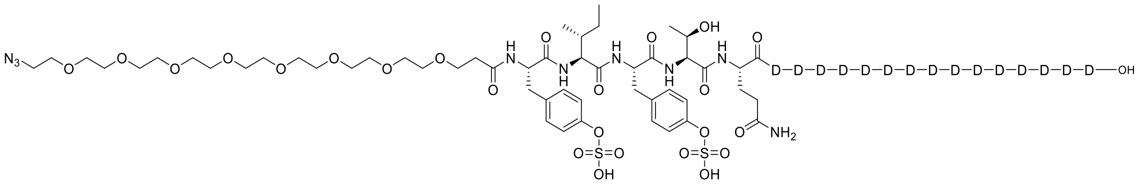


**B**

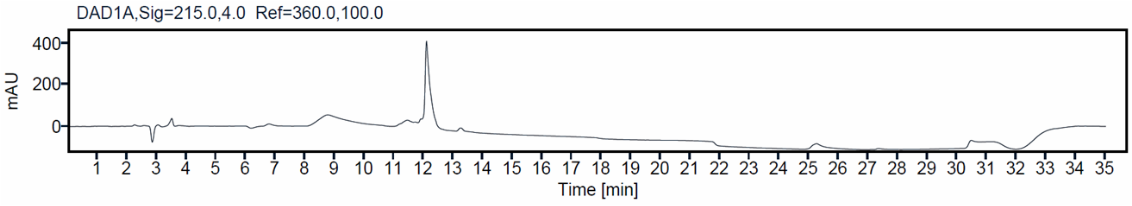


**C**

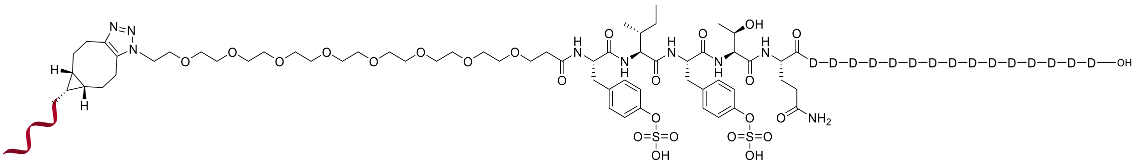


**D**

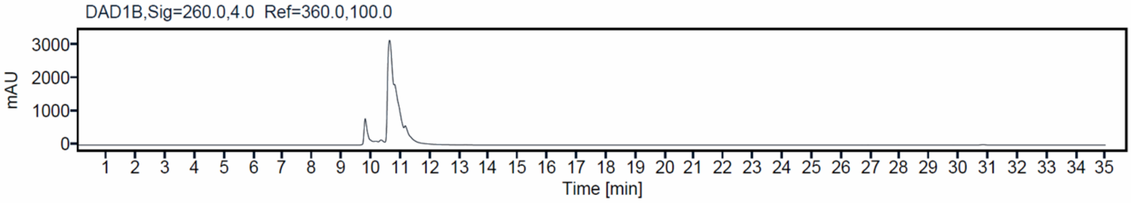


**E**

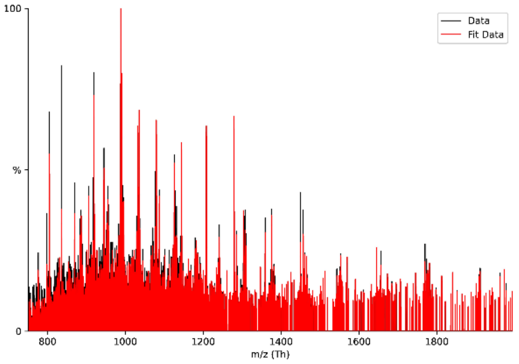


**F**

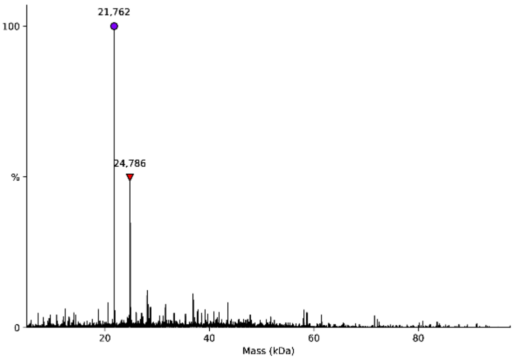


**Supplementary Figure S18.** (A) Chemical structure of azido-PEG$_8$-Tyr(SO$_3$H)-Ile-Tyr(SO$_3$H)-Thr-Gln-(Asp)$_{15}$-OH. (B) UHPLC trace, the $t_R$ of the product is 12.1 min. HRMS (ESI): *m/z* = $[M-2H]^{2-}$ calc for $C_{112}H_{154}N_{24}O_{70}S_2$ 1509.4340, found 1509.4375; *m/z* = $[M-3H]^{3-}$ calc for $C_{112}H_{153}N_{24}O_{70}S_2$ 1005.9536, found 1005.9575. (C) Chemical structure of Template DNA-PEG$_8$-Tyr(SO$_3$H)-Ile-Tyr(SO$_3$H)-Thr-Gln-(Asp)$_{15}$-OH. (D) UHPLC trace, the $t_R$ of the product is 10.8 min. (E) Multiply charged ion series of MS spectrum in negative mode. (F) Deconvoluted HESI mass spectrum. Expected mass 24785, observed 24786. Observed 21762 is from unreacted BCN-C$_6$-Template DNA.
